## Supplementary information for "Optochemical profiling of NMDA receptor molecular diversity at synaptic and extrasynaptic sites"

**This PDF file includes:**

Supporting Texts 1 to 3
Figures S1 to S11
Tables S1 to S5
SI References

### Supporting Information Text

**Text S1: NMR and mass-spectroscopy characterization of MASp**

NMR spectra $^1$H (500.16 MHz) and $^{13}$C (125.78 MHz) for *trans* and *cis* MASp were recorded on a 500 Bruker spectrometer equipped with a sensitivity-optimized measurement head (cryoprobe). Chemical shifts ($\delta$, ppm) are given with reference DMSO-*d*6 for $^1$H and $^{13}$C NMR, respectively: 2.50, 39.51. Signal multiplicity is described as follows: s (singlet), d (doublet), t (triplet), and m (multiplet). Broad signals are described as br. Coupling constants (*J*) are given in hertz. Molecule numbering is only related to atom assignment, which was established on the basis of $^{13}$C using $^1$H decoupled spectra as well as correlation spectroscopy, heteronuclear single quantum coherence, and heteronuclear multiple bond coherence.

Mass spectra were recorded on an Orbitrap Exactive (Thermoscientifc) mass spectrometer with positive (ESI+) electrospray ionization (ionization tension, 3.2 kV; ion transfer tube temperature, 275 °C). HPLC-MS analysis was performed on an Orbitrap Exactive Instrument as described above, equipped for HPLC (Nexera X2, Shimadzu) with a Phenomenex Kinetex C18 column (50 mm × 2.1 mm, 2.6 $\mu$m). MASp was eluted with the following gradient using solvent A ($H_2O/HCO_2H$: 100/0.1) and solvent B ($MeCN/HCO_2H$: 100/0.1), flow rate: 0.2 mL/min: 2% B linear increase to 100% B for 15 min, 100% B from 15 to 20 min, and linear decrease to 2% B from 20 to 20.01 min.

$^1$H NMR (500 MHz, DMSO-$d_6$) $\delta$: 9.16 (brs, 2H, $NH_2^+$), 8.93 (brs, 4H, $2NH_2^+$), 8.02 (brs, 3H, $NH_3^+$), 7.87 (d, $J_{H\text{-}10,H\text{-}11}$ = 9.0 Hz, 1H, H-10), 7.82 (d, $J_{H\text{-}7,H\text{-}6}$ = 9.0 Hz, 1H, H-7), 7.17 (d, $J_{H\text{-}11,H\text{-}10}$ = 9.0 Hz, 1H, H-11), 7.07 (d, $J_{H\text{-}6,H\text{-}7}$ = 9.0 Hz, 1H, H-6), 7.06 (s, 1H, H-2), 4.36 (t, $J_{H\text{-}13,H\text{-}14}$ = 5.0 Hz, 2H, H-13), 4.22 (t, $J_{H\text{-}4,H\text{-}3}$ = 5.5 Hz, 2H, H-4), 3.83 (t, $J_{H\text{-}3,H\text{-}4}$ = 5.5 Hz, 2H, H-3), 3.42 (t, $J_{H\text{-}14,H\text{-}13}$ = 5.0 Hz, 2H, H-14), 3.12 (t, $J_{H\text{-}24,H\text{-}23}$ = 7.5 Hz, 2H, H-24), 3.02 (t, $J_{H\text{-}22,H\text{-}23}$ = 7.5 Hz, 2H, H-22), 2.99 (t, $J_{H\text{-}15,H\text{-}16}$ = 7.5 Hz, 2H, H-15), 2.93 (m, 4H, H-18, H-21), 2.89 (t, $J_{H\text{-}17,H\text{-}16}$ = 7.5 Hz, 2H, H-17), 2.02 (tt, $J_{H\text{-}23,H\text{-}22}$ = $J_{H\text{-}23,H\text{-}24}$ = 7.5 Hz, 2H, H-23), 1.90 (tt, $J_{H\text{-}16,H\text{-}15}$ = $J_{H\text{-}16,H\text{-}17}$ = 7.5 Hz, 2H, H-16), 1.64 (m, 4H, H-19, H-20).

$^{13}$C NMR (126 MHz, DMSO-$d_6$) $\delta$: 170.9 (C-1), 160.3 (C-5), 159.9 (C-12), 158.7 (q, $^3J_{C,F}$ = 31.5 Hz, $CO_2^-$), 146.6 (C-9), 146.3 (C-8), 134.7 (C-2), 124.2 (C-7), 124.1 (C-10), 117.1 (q, $^1J_{C,F}$ = 299.5 Hz, C-F), 115.2 (C-11), 115.1 (C-6), 65.0 (C-4), 63.8 (C-13), 46.1 (C-18, C-21), 46.0 (C-14), 44.3 (C-24), 43.9, 43.8 (C-22, C-15), 36.6 (C-3), 36.2 (C-17), 23.8 (C-16), 22.7 (C-19, C-20), 22.4 (C-23).

HRMS (ESI+) m/z: calcd for $[C_{30}H_{43}N_7O_4 + H]^+$, 566.3449; found, 566.3436 (−2.2954 ppm); calcd for $[C_{30}H_{43}N_7O_4 + 2H]^{2+}$, 283.6761; found, 283.6754 (−2.4676 ppm); HPLC-MS (ESI) m/z (λ = 235 nm): RT = 6.76 min; 283.6755 $[M + 2H]^{2+}$ (−2.2637 ppm).

**Spectra S1: NMR characterization of MASp**

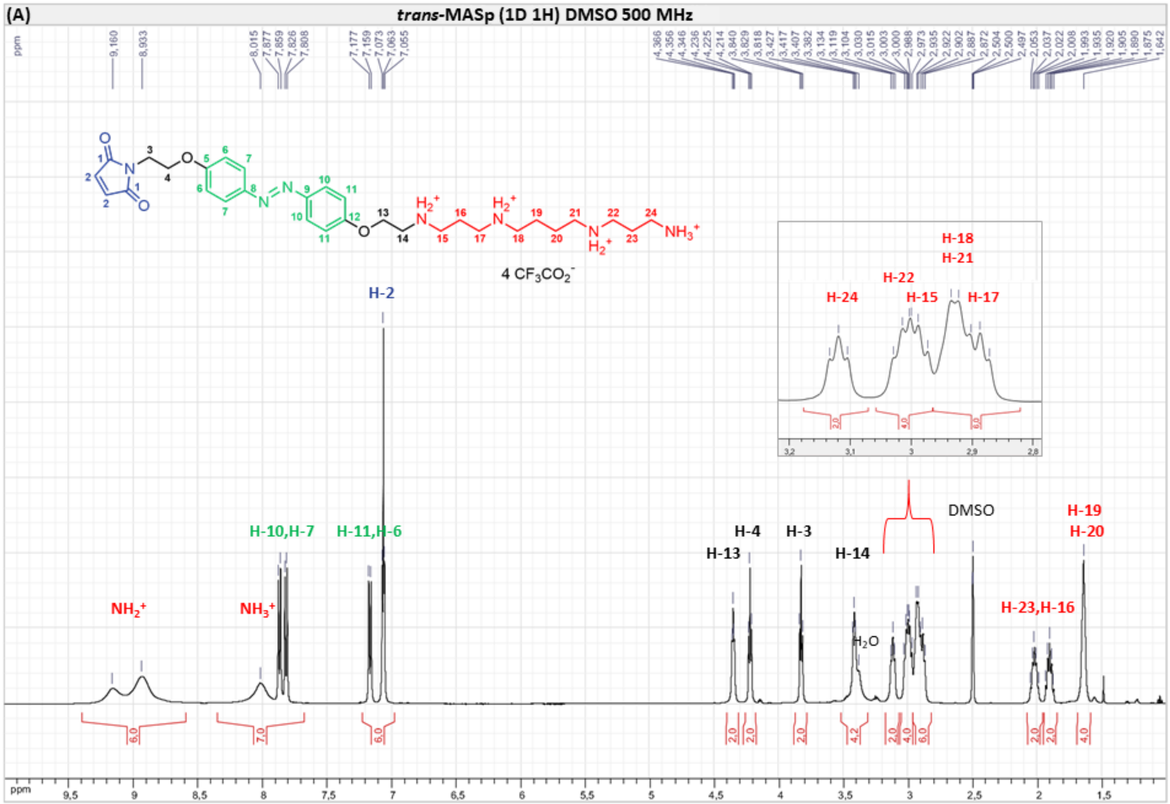


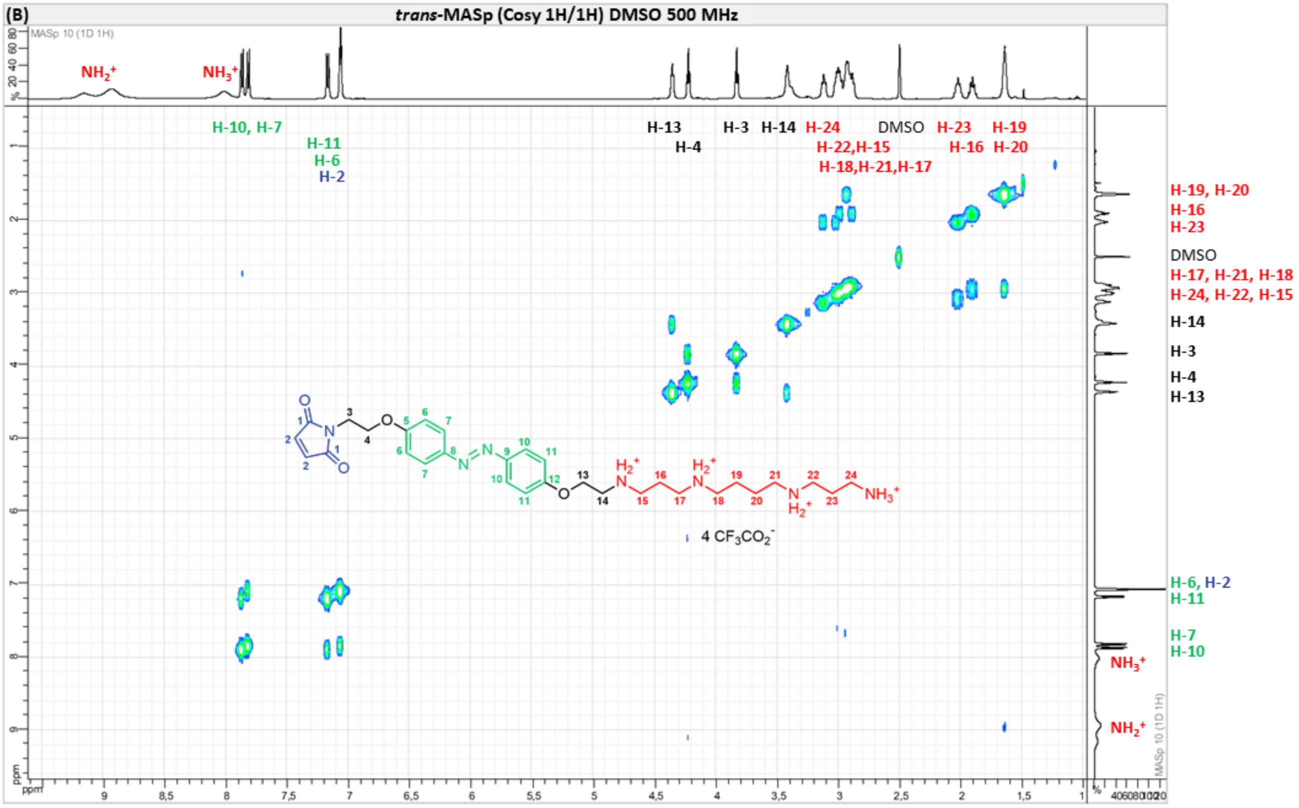

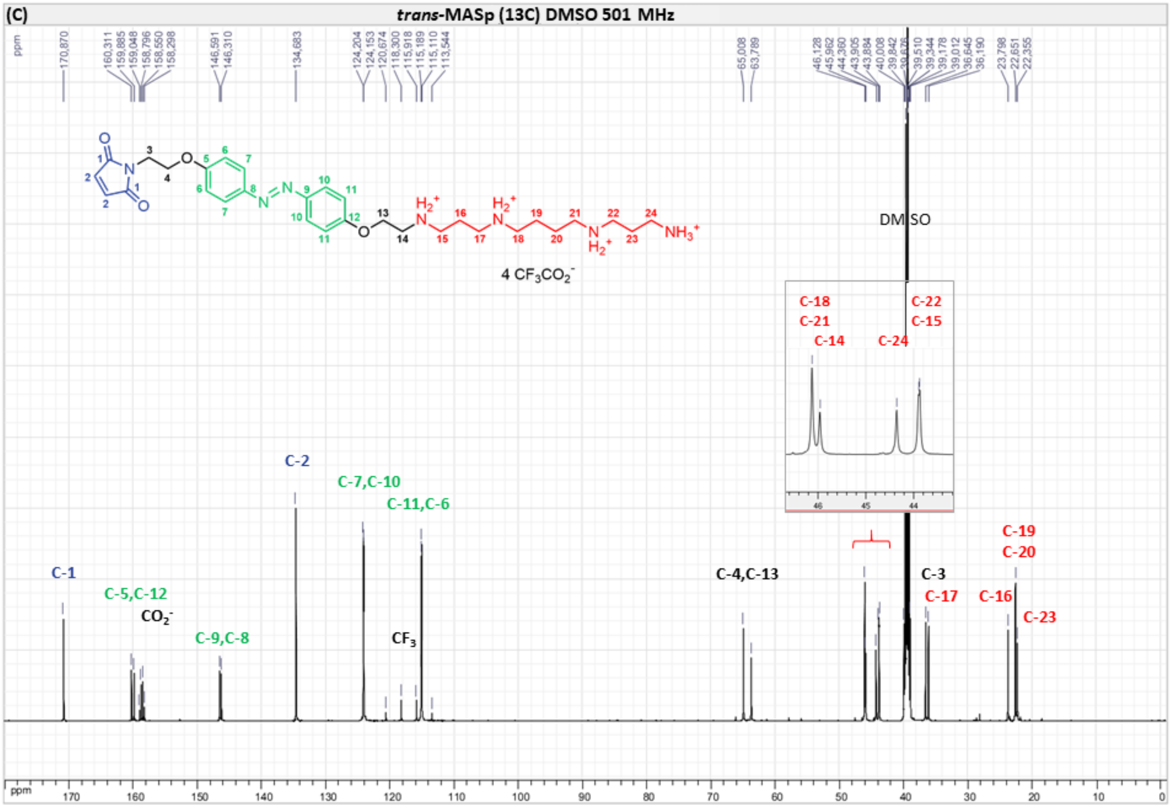
(C)
trans-MASp (13C) DMSO 501 MHz
170,870
160,311
159,885
159,048
158,796
158,550
158,298
146,591
146,310
134,683
124,204
124,153
120,674
118,300
115,918
115,189
115,110
113,544
65,008
63,789
46,128
45,962
44,360
43,905
43,884
40,008
39,842
39,676
39,510
39,344
39,178
39,012
36,645
36,190
23,798
22,651
22,355
4 CF3CO2-
DMSO
C-18
C-21
C-14
C-22
C-15
C-24
C-1
C-5,C-12
CO2-
C-9,C-8
C-2
C-7,C-10
C-11,C-6
CF3
C-4,C-13
C-3
C-17
C-16
C-19
C-20
C-23
ppm

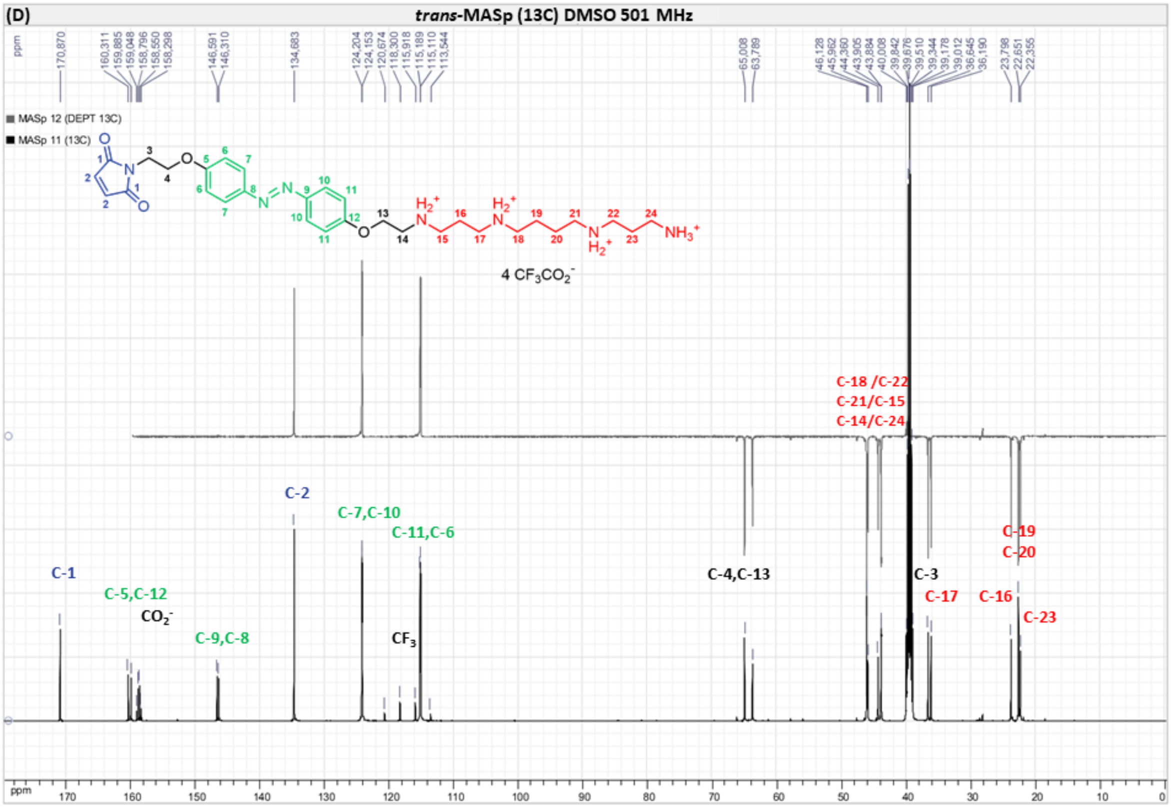
(D)
trans-MASp (13C) DMSO 501 MHz
MASp 12 (DEPT 13C)
MASp 11 (13C)
4 CF3CO2-
C-18 /C-22
C-21/C-15
C-14/C-24
C-1
C-5,C-12
CO2-
C-9,C-8
C-2
C-7,C-10
C-11,C-6
CF3
C-4,C-13
C-3
C-17
C-16
C-19
C-20
C-23
ppm

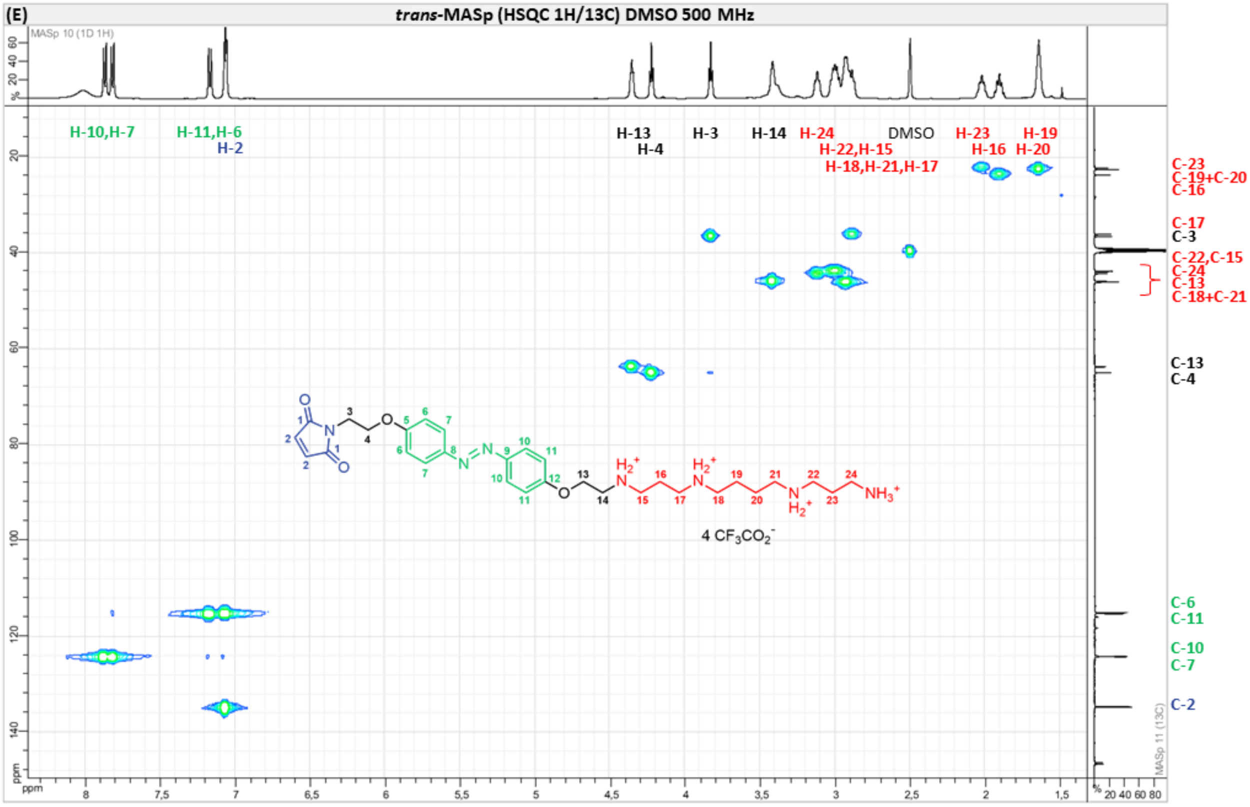
(E)
trans-MASp (HSQC 1H/13C) DMSO 500 MHz
MASp 10 (1D 1H)
H-10,H-7
H-11,H-6
H-2
H-13
H-4
H-3
H-14
H-24
H-22,H-15
H-18,H-21,H-17
DMSO
H-23
H-16
H-19
H-20
C-23
C-19+C-20
C-16
C-17
C-3
C-22,C-15
C-24
C-13
C-18+C-21
C-13
C-4
C-6
C-11
C-10
C-7
C-2
4 CF3CO2-
MASp 11 (13C)
ppm

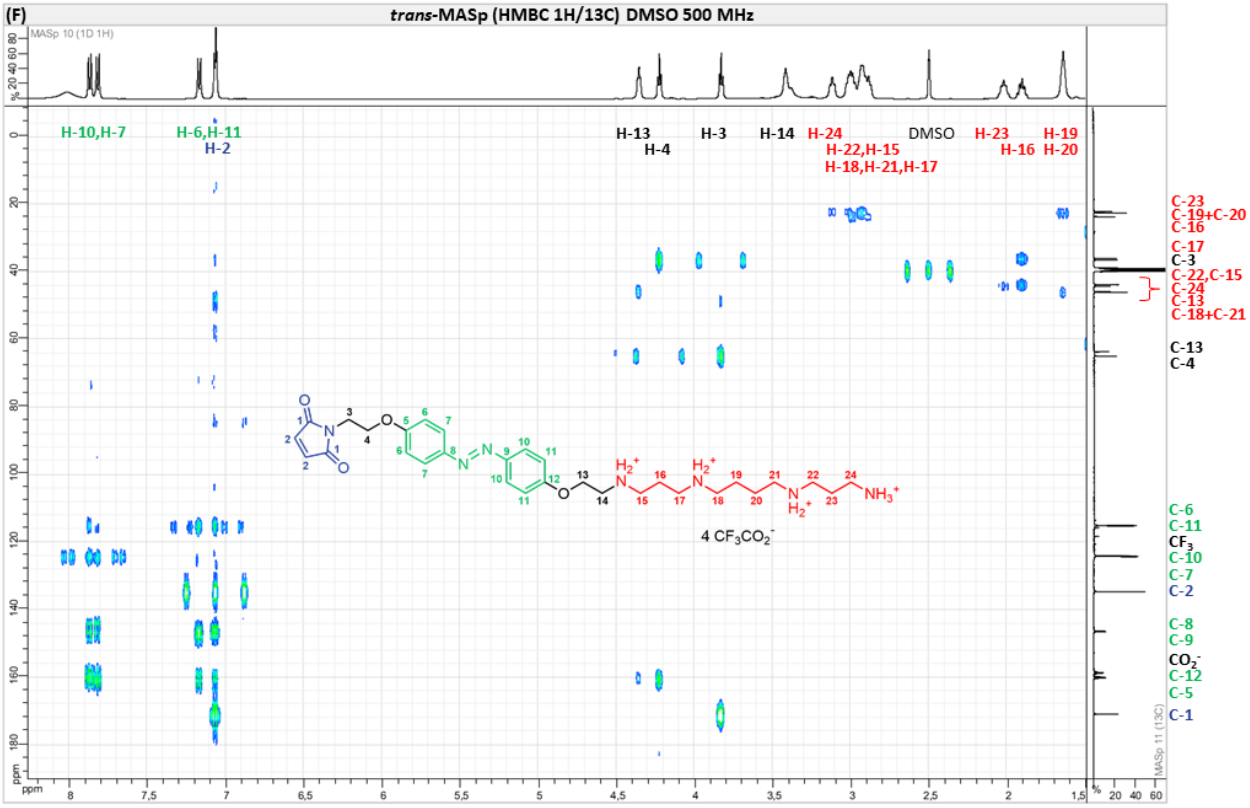
(F)
trans-MASp (HMBC 1H/13C) DMSO 500 MHz
MASp 10 (1D 1H)
H-10,H-7
H-6,H-11
H-2
H-13
H-4
H-3
H-14
H-24
H-22,H-15
H-18,H-21,H-17
DMSO
H-23
H-16
H-19
H-20
C-23
C-19+C-20
C-16
C-17
C-3
C-22,C-15
C-24
C-13
C-18+C-21
C-13
C-4
C-6
C-11
CF3
C-10
C-7
C-2
C-8
C-9
CO2-
C-12
C-5
C-1
4 CF3CO2-
MASp 11 (13C)
ppm

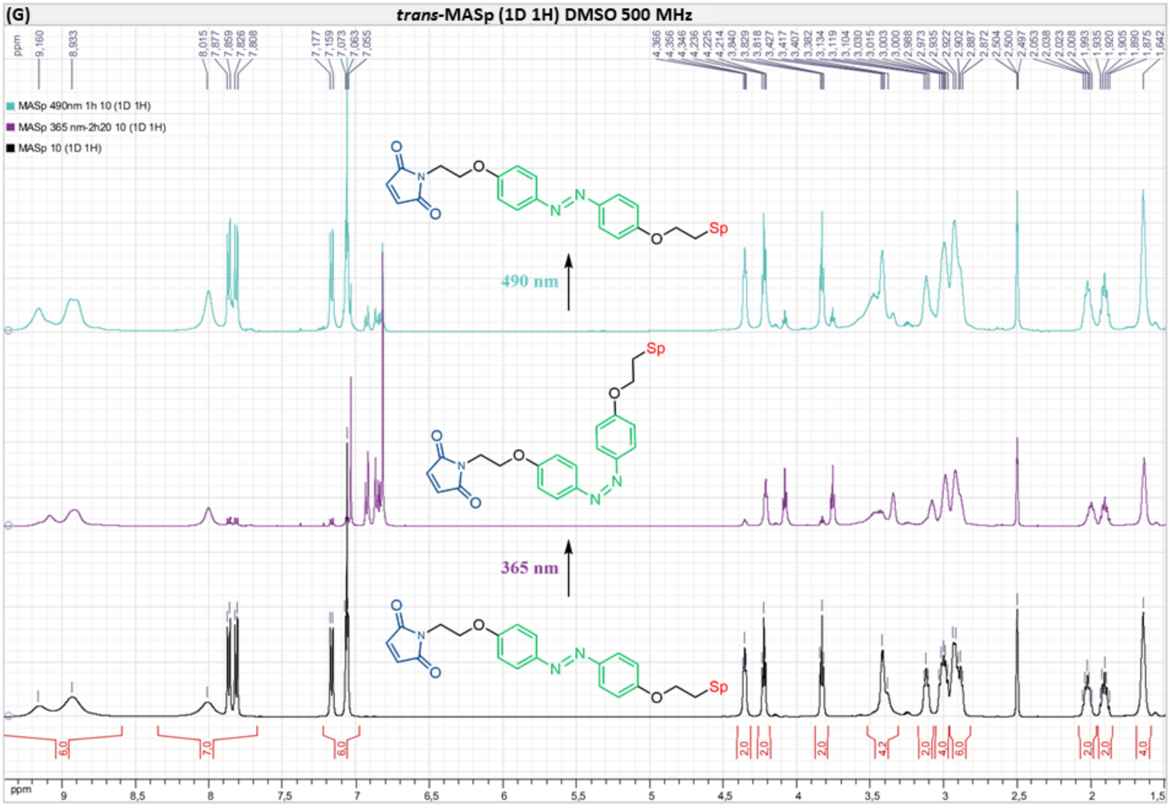


Lower panel: $^1$H NMR spectrum of MASp in the dark (100% *trans*). Middle panel: $^1$H NMR spectrum of MASp 365 nm PSS. Upper panel: $^1$H NMR spectrum of MASp 490 nm PSS.

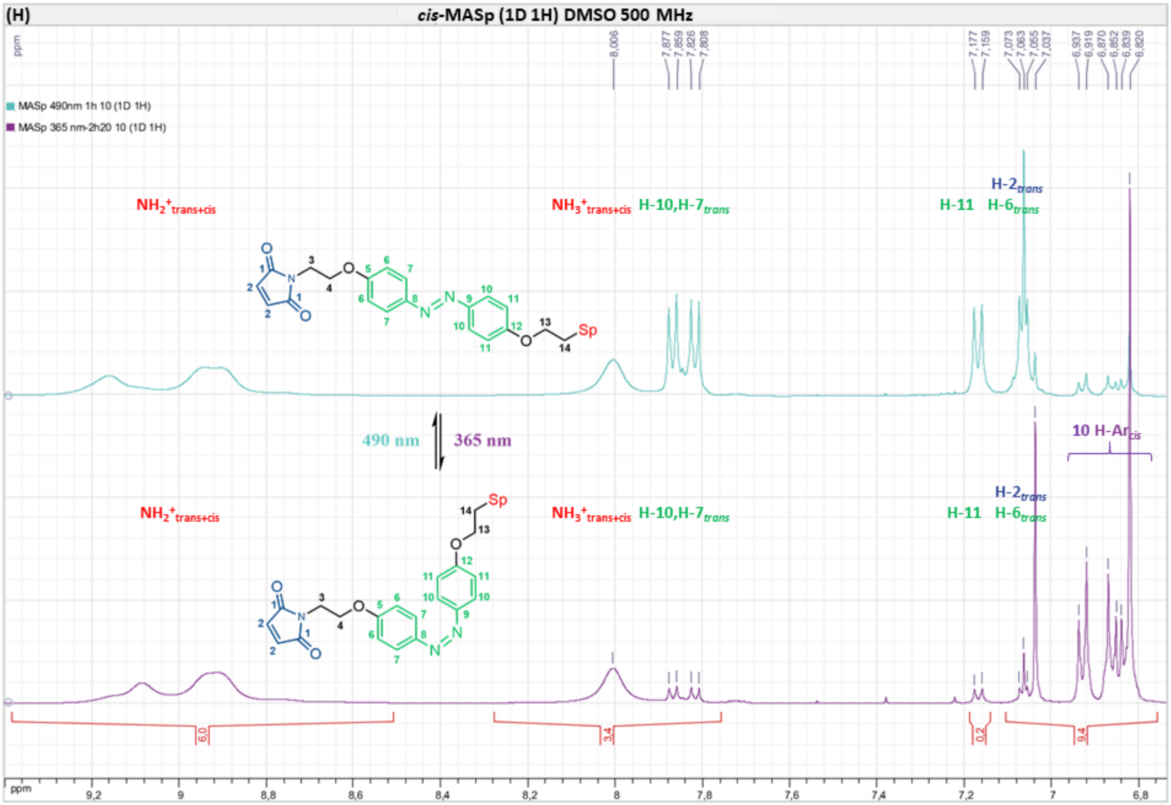


Percentage of *cis* and *trans* isomers PSSs was calculated from the integration of the peaks at 8.20−7.80 ppm and 7.16 ppm. Lower panel: 90% *cis* and 10% *trans*. Upper panel: 91% *trans* and 9% *cis*.

**Spectra S2: Mass spectrometry characterization of MASp**

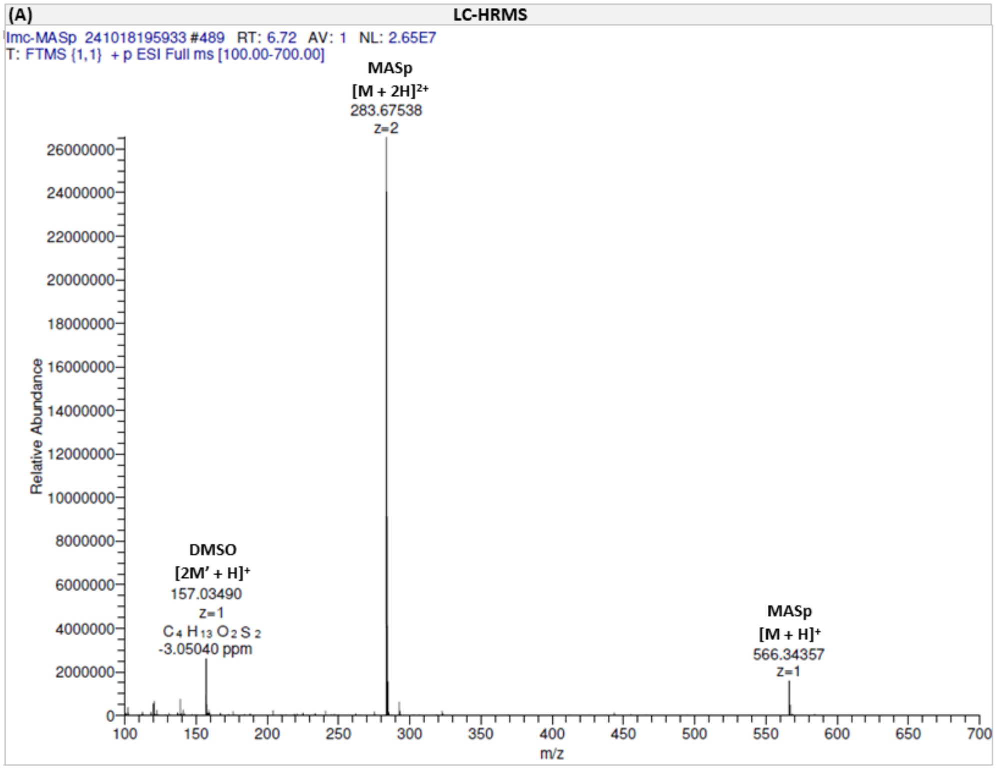


Full-MS spectrum at 6.72 min retention time showing essentially z = 2.

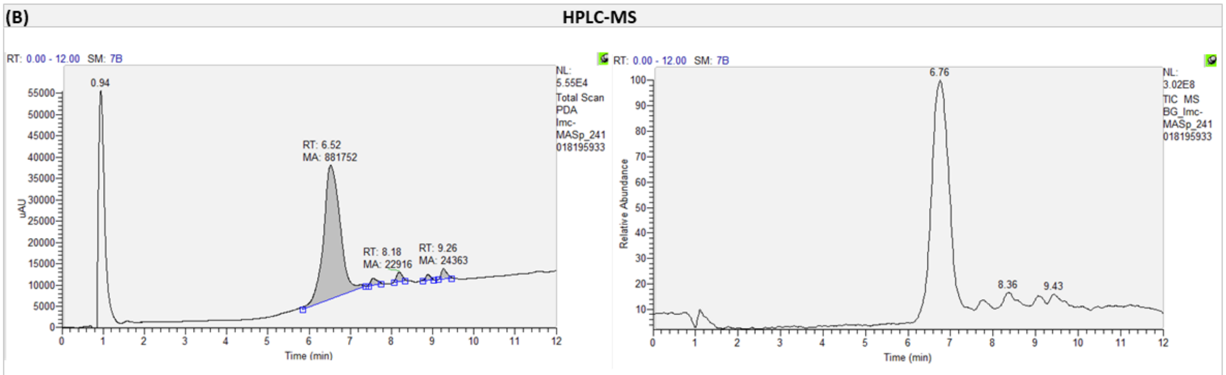


| PEAK LIST | | |
|---|---|---|
| Imc-MASp_241018195933.raw | | |
| RT: 0.00 - 12.00 | | |
| Number of detected peaks: 5 | | |
| Apex RT | Area | %Area |
| 6.52 | 881751.744 | 91.66 |
| 7.55 | 19748.257 | 2.05 |
| 8.18 | 22916.46 | 2.38 |
| 8.88 | 13154.794 | 1.37 |
| 9.26 | 24362.606 | 2.53 |

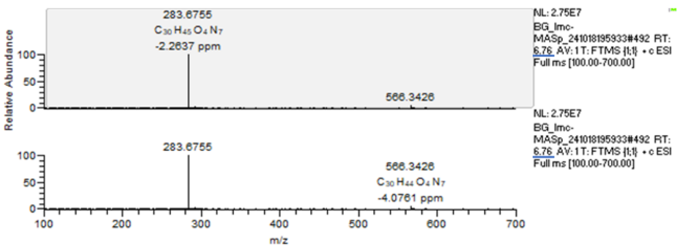


Left panel: UV detection chromatogram (PDA). Right panel: Mass spectrum peaks (TIC).

**Text S2 (related to Fig. S3): Photoswitching kinetics in mammalian cells**

At the maximal light power in our experimental conditions (~10 mW for 365 nm and 27 mW for 525 nm), receptor activity could be potentiated by UV light with a time constant of ~60 ms (MASp *trans*-to-*cis* transition) and reverted back to its basal state by 525 nm light with a much slower time constant of ~450 ms (MASp *cis*-to-*trans* transition; Fig. S3E,F). Strikingly, we observed that using blue light (460 nm) instead of green light (525 nm) to induce MASp *trans*-to-*cis* transition allowed a much faster (30-fold) reversal of receptor potentiation (time constant of ~13 ms; Fig. S3E,F), indicating that the slow transition under 525 nm light does not reflect slow kinetics of receptor conformational changes. Current off-rates with 525 nm light were systematically slower than current off-rates with 460 nm light regardless of the light power used (Fig. S3G). Hence, the wavelength-dependence of receptor off-rate kinetics is likely due to the much smaller absorbance of *cis*-MASp at 525 compared to 460 nm (*cis*-MASp absorptivity ε ~ 1900 $L.mol^{-1}.cm^{-1}$ at 460 nm and ε ~ 200 $L.mol^{-1}.cm^{-1}$ at 525 nm; calculated from the 365 nm PSS spectrum of Fig. S1A), which results in a smaller probability of the molecule to undergo the *cis*-to-*tran*s transition at 525 nm relative to 460 nm. Return to basal activity was however less complete under 460 nm light compared to 525 nm light (Fig. S3E,F), consistently with the lower proportion of the *trans* isomer in the 460 nm PSS compared to the 525 nm PSS (as assessed by UV-visible spectroscopy; Fig. S1A). MASp thus allows very fast modification of GluN1/GluN2B diheteromer activity with on and off kinetics that can be below 100 ms. Depending on the requirements of the biological experiment, reversal of receptor potentiation can be performed either with blue light to achieve very fast reversal, or with green light to achieve more complete reversal to the basal state at the cost of a slower recovery. In the paper we will favor the second option and use 365 nm and 525-530 nm as the two photoswitching wavelengths.

**Text S3 (related to Fig. S7). Agonist-dependence of UV-induced potentiation**

To quantify the amount of photomodulation of tonic currents by UV light, we first assessed the glutamate-dependence of the photomodulation ratio when measured at the peak or steady-state UV-potentiation. For this we turned to Xenopus oocytes in order to obtain expression of a pure population of GluN2B*-R187C diheteromers. Similarly to neurons, we observed on this system sustained UV potentiation at saturating glutamate concentrations (100 µM) and a transient potentiation at glutamate concentrations lower than the receptor $EC_{50}$ (0.03 and 0.1 µM; Fig. S7C,D). This was reflected in a decrease of glutamate $EC_{50}$ under UV light compared to under green light (Fig. S7E, see also Fig. S2E). As expected, the photomodulation ratio measured at the UV steady-state current ($I_{UV, SS}$ / $I_{490\ nm}$) displayed a clear glutamate-dependence (Fig. S7D). On the contrary, the photomodulation ratio measured at the UV peak current

($I_{UV, peak}$ / $I_{490\ nm}$) was nearly independent of glutamate concentration, with less than 10% difference of photomodulation ratios between the four glutamate concentrations tested (Fig. S7D).

We next investigated the glycine-dependence of photomodulation in presence of a saturating concentration of glutamate. Similarly to glutamate, UV induced a sustained potentiation for saturating (100 µM) and near-saturating (> 1 µM) concentrations of glycine but became transient at a glycine concentration lower than the $EC_{50}$ (0.1 µM; Fig. S7F,G). Current relaxation was however much less marked than for glutamate, with a steady-state current under UV light representing 72% of the peak current for 0.1 µM glycine (Fig. S7H) compared to 50% of the peak current for 0.1 µM glutamate (Fig. S7E). This decrease in UV steady-state at low glycine concentrations was not translated into a significant change of glycine $EC_{50}$ under UV light compared to 490 nm light (Fig. S7H). The photomodulation ratio measured at the UV peak slightly increased (by 20%) at low concentrations of glycine (Fig. S7G). In our *ex vivo* experiments, slices were perfused with 20 µM glycine so we can assume to be in a range of glycine concentrations for which the photomodulation ratio at the UV peak is constant.

### Supplementary Figures

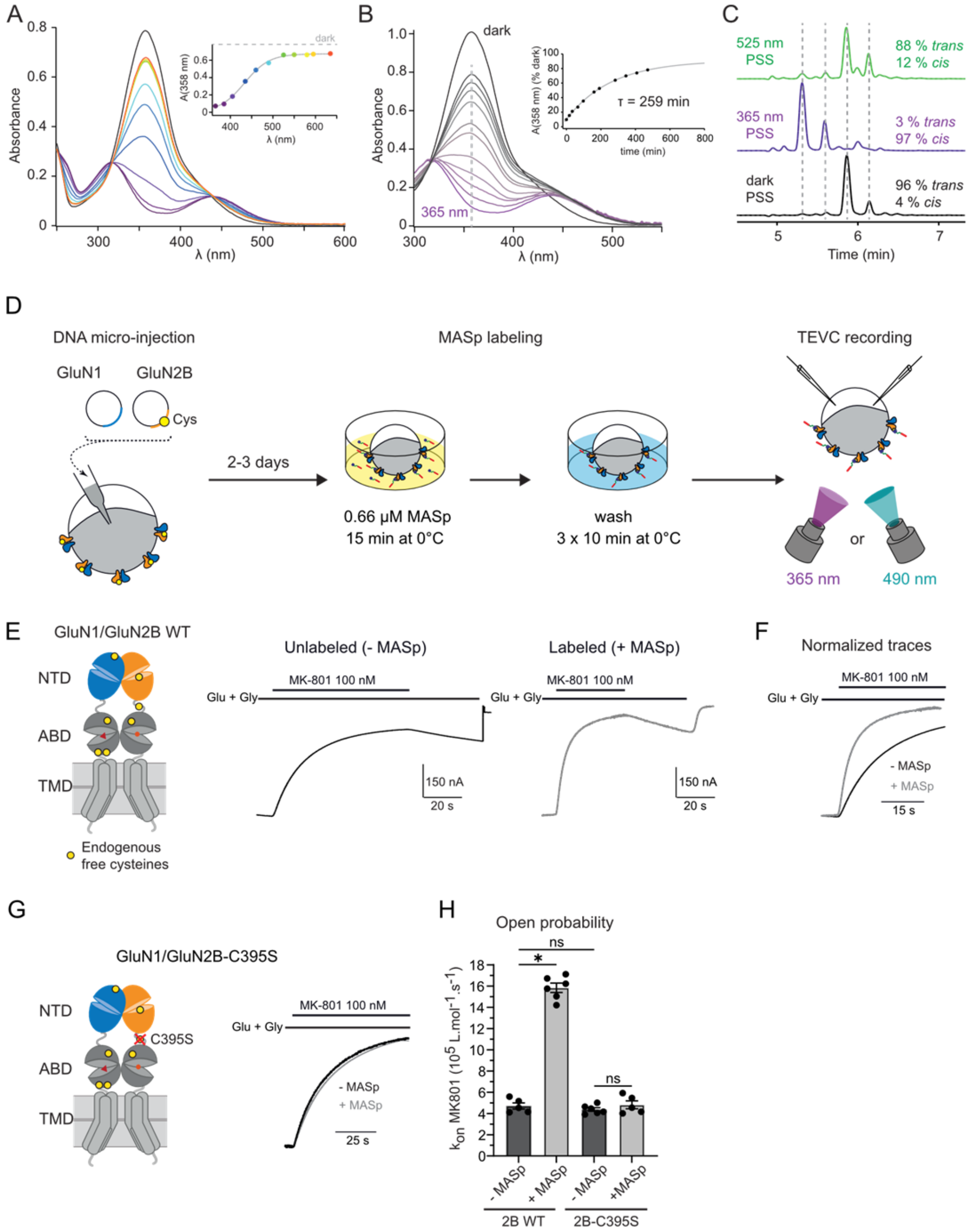


**Fig. S1 (related to Fig. 1): Photochemical properties of MASp and design of the labeling conditions in Xenopus oocytes.**

**(A)** UV-visible spectra of MASp conjugated to L-cysteine ($MASp_{Cys}$, see Methods and Fig. S11) in oocyte recording medium (pH 7.3, 33 μM MASp) in the dark (black trace) and under illumination with increasing wavelengths, from 365 nm (dark violet) to 635 nm (red). Inset, absorbance at 358 nm (*trans*-$MASp_{Cys}$ absorbance peak) as a function of the illumination wavelength. **(B)** Thermal stability of *cis*-$MASp_{Cys}$. After recording a UV-visible spectrum of $MASp_{Cys}$ in the dark (mainly *trans* state, black trace), $MASp_{Cys}$ was irradiated for 10 min by 365 nm light to yield mainly *cis*-$MASp_{Cys}$ (violet spectrum). Spectra in violet/grey gradation represent the gradual *cis*-to-*trans* transition of $MASp_{Cys}$ in the dark at different time points (up to ~8 h) post UV irradiation. Inset, evolution over time of the absorbance at 358 nm. Single-exponential fit (grey line) yielded a time constant $\tau$ = 259 min (~4 h) for *cis*-to-*trans* $MASp_{Cys}$ thermal relaxation in the dark. **(C)** HPLC chromatograms monitored at the isosbestic point (440 nm) of the photostationary states (PSS) of $MASp_{Cys}$ in the dark (black trace), after illumination with 365 nm light (violet trace) and subsequent illumination with 525 nm light (green trace). **(D)** Experimental workflow for heterologous expression of GluN1/GluN2B receptors, MASp labeling and photomodulation of NMDAR activity in Xenopus oocytes. **(E-H)** MASP binds to the endogenous cysteine GluN2B-C395. **(E)** Left, schematic of a GluN1/GluN2B dimer with the positions of the free cysteines (i.e. not involved in disulfide bridges) highlighted in yellow. Right, inhibition traces by MK-801 (100 nM), an open channel pore blocker, of unlabeled (- MASp, black) and labeled (+ MASp, grey) GluN1/GluN2B WT NMDARs kept in the dark (MASp in its *trans* state). **(F)** Superposition of the normalized MK801 inhibition traces from Panel (e). Note the increase of MK-801 inhibition rate after labeling with MASp, indicating an increase in the open probability when GluN1/GluN2B NMDARs are conjugated with MASp. **(G)** Left, neutralization of the reactivity of C395, a cysteine located in GluN2B NTD-ABD linker, by mutation into a serine. Right, superposed and normalized MK-801 inhibition traces for labeled (+ MASp, grey) and unlabeled (- MASp, black) GluN1/GluN2B-C395S mutants. Note that on this mutant, labeling does not affect the rate of inhibition by MK-801. **(H)** Summary of the rates of inhibition by MK-801. n.s., $p > 0.05$; *, $p < 0.05$; multiple Mann-Whitney tests, p-values were adjusted for multiple comparisons using Bonferroni correction. Only the pre-selected indicated comparisons were performed.

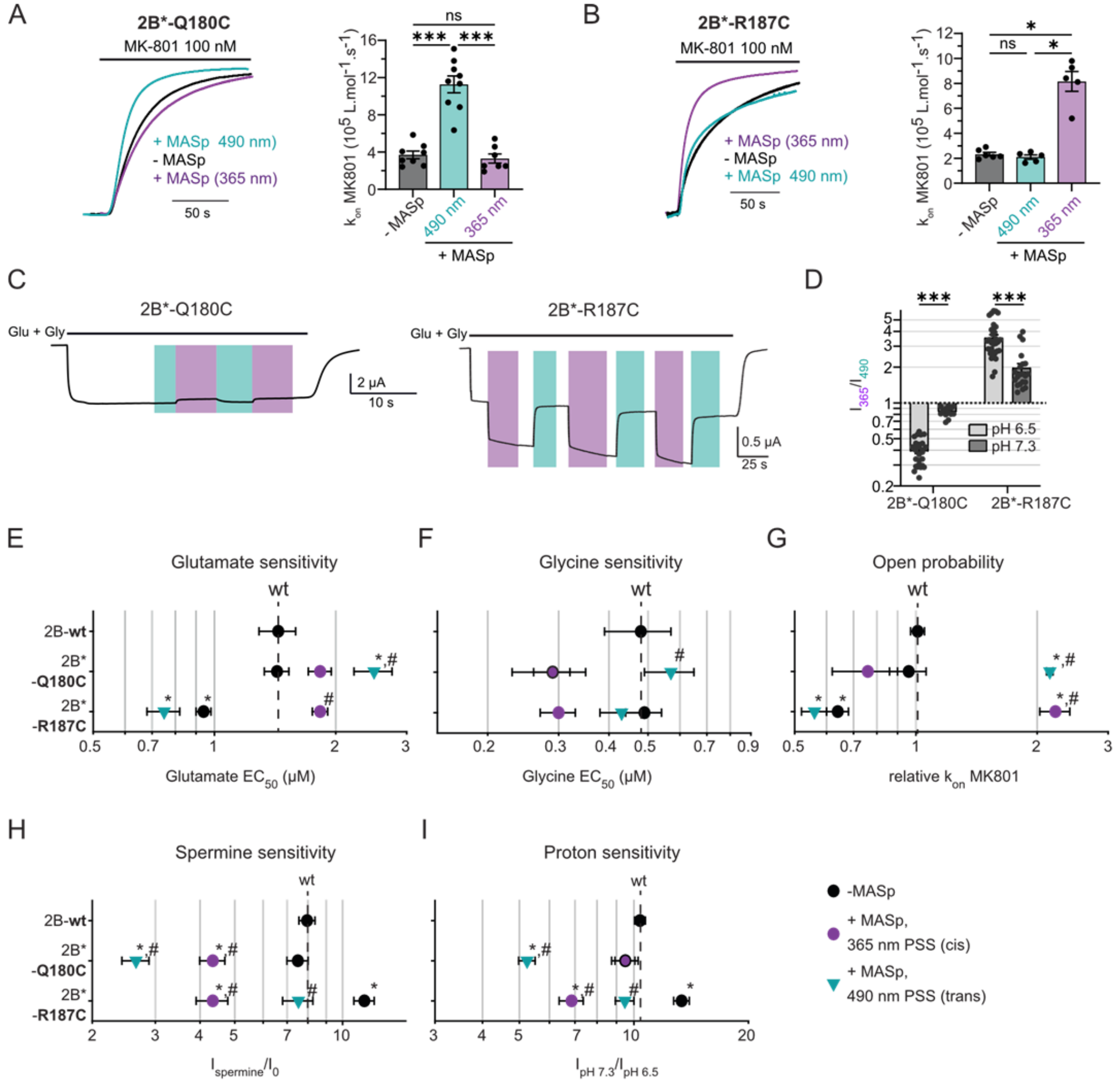


J

| Criteria | Q180C | R187C |
|---|---|---|
| Photomodulation at physiological pH | 1.2-fold | 2-fold |
| *cis*-active photoswitch? | ✗ | ✓ |
| Functionally silent cysteine mutation | ✓ | ✓ |
| Photoswitch inert in one configuration | ✓ | ✓ |

**Fig. S2 (related to Fig. 1): Photomodulation and pharmacological properties of GluN2B*-Q180C and -R187C mutants.**

**(A,B)** Evaluation of the relative Po of MASp-labeled GluN1/GluN2B*-Q180C (**A**) and GluN1/GluN2B*-R187C (**B**) under different light conditions. Left, superposed current traces following application of agonists and 100 nM MK-801 of unlabeled receptors (- MASp, black) and MASp-labeled receptors (+ MASp) under 365 (violet, mostly *cis*-MASP) or 490 nm light (blue-green, mostly *trans*-MASp). Right, summary of MK-801 inhibition rates. Kinetics of MK801 inhibition: 2B*-Q180C, $k_{on}$ = 3.7 ± 0.4 $10^5$ $L.mol^{-1}.s^{-1}$, n = 8 (- MASp); 11.3 ± 0.9 $10^5$ $L.mol^{-1}.s^{-1}$, n = 9 (+ MASp, 490 nm), and 3.3 ± 0.5 $10^5$ $L.mol^{-1}.s^{-1}$, n = 7 (+ MASp, 365 nm); 2B*-R187C, $k_{on}$ = 2.3 ± 0.1 $10^5$ $L.mol^{-1}.s^{-1}$, n = 6 (- MASp); 2.1 ± 0.2 $10^5$ $L.mol^{-1}.s^{-1}$, n = 5 (+ MASp, 490 nm), and 8.2 ± 1.6 $10^5$ $L.mol^{-1}.s^{-1}$, n = 5 (+ MASp, 365 nm). n.s., $p > 0.05$; *, $p < 0.05$; ***, $p < 0.001$; multiple Mann-Whitney test, p-values were adjusted for multiple comparisons using Bonferroni correction. When MASp was conjugated to GluN2B-Q180C in its *trans* configuration (under 490 nm light), channel Po increased by ~2-fold compared to unlabeled receptors, while labeling with *cis*-MASp (under UV light) produced no significant change in receptor activity. MASp thus acts as a *trans*-on photoswitch at position Q180C. In contrast, MASp conjugated to GluN2B-R187C had no effect on receptor channel Po under 490 nm light, while it increased Po by ~3.5-fold under UV-light. MASp thus acts as a *cis*-on photoswitch at position R187C (see Fig. 1E, right). **(C,D)** Photomodulation of MASp-labeled, GluN1/GluN2B*-Q180C and -R187C receptors at physiological pH. **(C)** Current traces from MASp-labeled GluN1/GluN2B*-Q180C (left) and GluN1/GluN2B*-R187C (right) at pH 7.3. **(D)** Summary of photomodulation ratios for the Q180C and R187C mutants at pH 6.5 and 7.3. Photomodulation values at pH 6.5 are from Fig. 1. ***, $p < 0.001$ multiple Mann-Whitney tests, p-values were adjusted for multiple comparisons using Bonferroni correction. Only the pre-selected indicated comparisons were performed. Photomodulation values and number of cells are summarized in Table S1. **(E-I)** Determination of the relative open probability and sensitivity to different pharmacological agents of unlabeled (- MASp) and labeled (+ MASp) GluN1/GluN2B*-Q180C and GluN1/GluN2B*-R187C receptors compared to unlabeled WT GluN1/GluN2B receptors. *, $p < 0.05$ between the condition and unlabeled WT GluN1/GluN2B receptors; #, $p < 0.05$, between MASp-labeled mutant receptors under 365 or 490 nm light and unlabeled mutant receptors. One way Anova followed by Tukey's test. **(J)** Pros and cons of photo-enhancing GluN2B-NMDARs via the Q180C or the R187C labeling position. The R187C is the most favored labeling position due to its strong photomodulation at physiological pH and the fact that MASp at this position is active in *cis*.

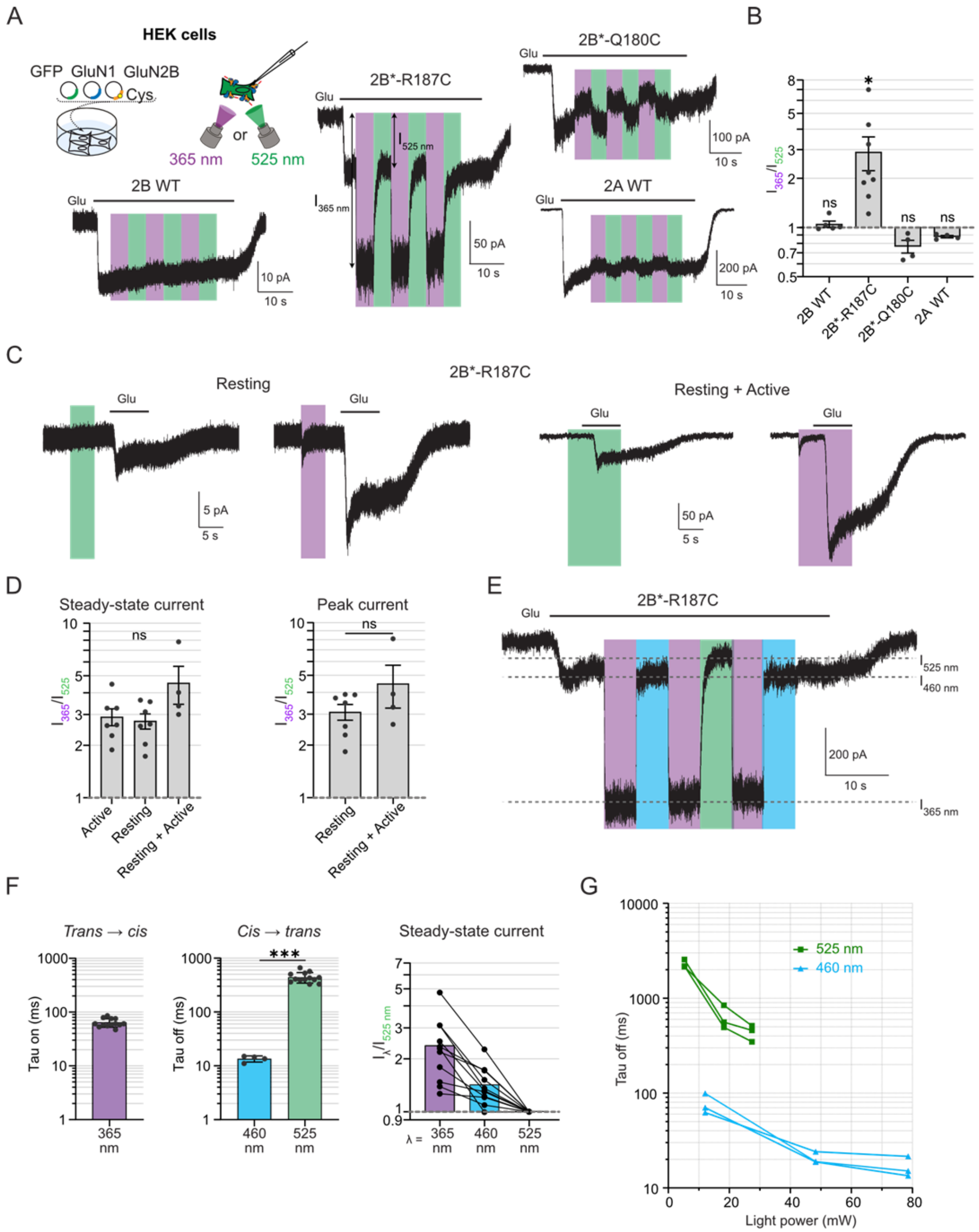


**Fig. S3 (related to Fig. 2): Up to 3-fold potentiation of GluN2B diheteromer currents in HEK cells.**

**(A)** Current traces from MASp-labeled HEK cells expressing GluN1/GluN2B WT, GluN1/GluN2B*-Q180C, GluN1/GluN2B*-R187C, and GluN1/GluN2A WT receptors, following application of glutamate (100 µM) under 365 nm (violet bars) or 525 nm (green bars) illumination. Cells were constantly perfused with glycine (100 µM). **(B)** Summary of the photomodulation ratios ($I_{365\,nm}$ / $I_{525\,nm}$) of currents from the different receptors in (a) expressed in HEK cells. n.s., $p > 0.05$; **, $p < 0.01$; multiple one sample Wilcoxon tests against the value 1, p-values were adjusted for multiple comparisons using Bonferroni correction. Photomodulation values and number of cells are indicated in Table S1. **(C,D)** No state-dependence of MASp-induced photomodulation. **(C)** GluN1/GluN2B*-R187C current traces in HEK cells under different illumination protocols. Left, light was applied before agonist application, allowing receptor modulation only from its resting state (Resting). Right, light was applied before and during agonist application, allowing receptor modulation from both its resting and activated states (Resting + Active). The small current peak observed at the onset of UV light in absence of agonist application corresponds to a transient potentiation of NMDARs tonically activated by small amounts of contaminating glutamate (see Fig. S7). **(D)** Photomodulation ratios of steady-state (left) and peak (right) currents following light application during the active state only (Active, panel a), the resting state only (Resting, panel c, left) and both the active and resting states (resting + Active, panel c, right). Photomodulation ratios under these three states were measured from HEK cells from the same coverslip (hence labeled at the same time) to minimize any variability linked to the labeling procedure. n.s., $p > 0.05$, Kruskall Wallis (left) and Mann Whitney (right) tests. **(E-G)** Wavelength-dependence of MASp-induced photomodulation. **(E)** GluN1/GluN2B*-R187C current trace in a MASp-labeled HEK cell following illumination with UV (365 nm, violet bar), blue (460 nm, blue bar) or green (525 nm, green bar) light. Cells were pre-illuminated with blue light before current recording. **(F)** Time constants of current potentiation by UV light (left) and current recovery from potentiation by blue or green light (middle), and steady-state currents under the different light conditions relative to the current under green light, which represents the receptor basal activated state (right). Kinetics of photomodulation were measured at maximal light power through a 10x objective (~10 mW for 365 nm, 79 mW for 460 nm and 27 mW for 525 nm). Tau on (365 nm) = 64 ± 13 ms (n = 13), Tau off (460 nm) = 13.5 ± 0.9 ms (n = 4), Tau off (525 nm) = 442 ± 27 ms (n = 13). ***, $p < 0.001$, Mann Whitney test. Relative steady-state (SS) values: SS(365 nm) = 2.38 ± 0.30; SS(460 nm) = 1.43 ± 0.11. **(G)** Kinetics of current recovery from potentiation by blue or green light as a function of light power. Recordings in HEK cells were performed at pH 7.3, -60 mV.

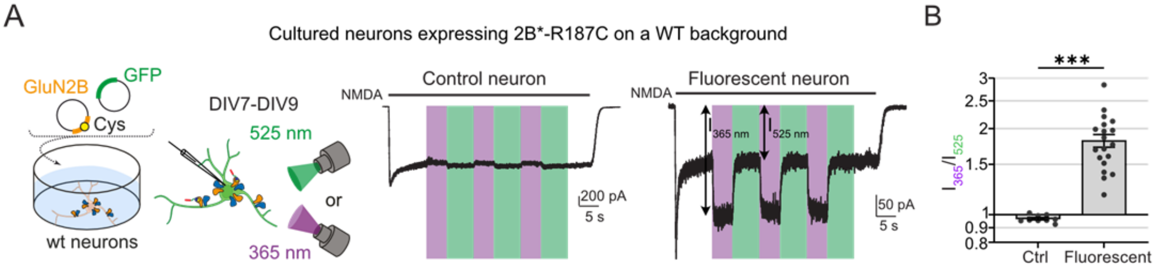


**Fig. S4 (related to Fig. 2): Strong photomodulation of GluN2B*-R187C diheteromers in cultured cortical neurons.**

**(A)** Left, expression of the GluN2B*-R187C subunit in cultured cortical neurons from wild-type animals through *ex utero* electroporation (see Methods). Right, current traces from DIV7-9 neurons labeled with MASp, following application of the selective NMDAR agonist NMDA (300 µM), under 365 nm (violet bars) or 525 nm (green bars) illumination. Cells were constantly perfused with D-serine (50 µM). Fluorescent neurons express the 2B*-R187C subunit, while non-fluorescent (control) neurons only express the endogenous NMDAR population. (**B**) Summary of photomodulation ratios of MASp-labeled, fluorescent and control (non-fluorescent) neurons. ***, $p < 0.001$, Mann Whitney test. Recordings in cultured cortical neurons were performed at pH 7.3, -60 mV. Photomodulation values and number of cells are summarized in Table S1.

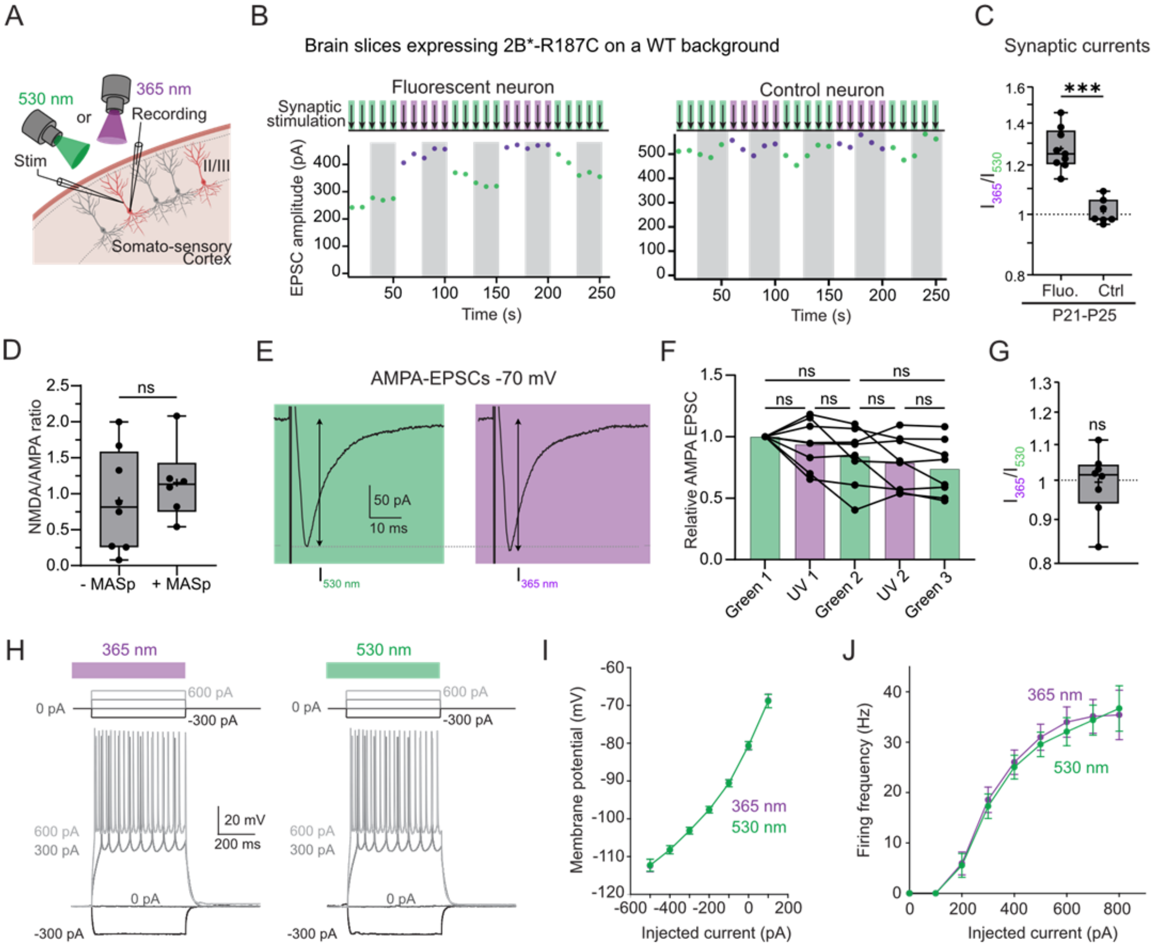


**Fig. S5 (related to Fig. 3): Protocol of photomodulation of NMDA-EPSCs and determination of MASp background effects on brain slices.**

**(A)** Data were obtained from electroporated (fluorescent) and non-electroporated (non fluorescent, control) layer II/II pyramidal neurons from the somatosensory cortex upon stimulation at the level of their apical dendrite. **(B)** Top, protocol of photomodulation of NMDAR synaptic currents (NMDA-EPSCs). Synaptic stimulation was performed continuously at 0.1 Hz. 525 nm (green bars) or 365 nm (violet bars) illumination started 1 s before synaptic stimulation and ended 200 ms after synaptic stimulation (1.2 s total). Illumination cycles consisted of alternations between 5 stimulations under 525 nm light and 5 stimulations under 365 nm light. Bottom, amplitudes of NMDAR-EPSCs from a MASp-labeled, fluorescent (expressing GluN2B*-R187C) neuron from a P14 mouse under alternating 525 nm (green points) and 365 nm (violet points) illumination, showing reversible and reproducible photomodulation of NMDA-EPSCs. Given the slow reversion of EPSC amplitude by 525 nm light, only the three last EPSCs of each 5-stimulation cycle were considered for amplitude quantification and trace averaging (grey bars). **(c)** Photomodulation ratios ($I_{365\ nm}$ / $I_{525\ nm}$) of NMDA-EPSCs of MASp-labeled, fluorescent and control neurons from P21-P25 animals. Photomodulation values and number of cells are summarized in

Table S3**. (D-J)** MASp labeling has no or minimal impact on synaptic transmission and neuronal electrical properties. **(D)** NMDA/AMPA ratios for WT neurons either unlabeled (- MASp, n = 8) or labeled (+ MASp, n = 6) from P21-P25 animals. n.s., $p > 0.05$, Mann Whitney test. **(E-G)** No photodependent effect of MASp labeling on AMPA-EPSCs. **(E)** Representative AMPA-EPSCs (recorded at -70 mV) from a MASp-labeled control (non-fluorescent) layer II/III cortical neuron under 525 nm (green bar, left) or 365 nm light (violet bar, right) in a slice from a P21 mouse. Same illumination conditions as in Panel B. **(F)** Summary of AMPA-EPSC amplitudes of MASp-labeled control neurons from P21-P22 mice under alternating 525 nm (green bars) and 365 nm (violet bars) illumination. Each dot represents the average amplitude of AMPA-EPSCs for each cycle for each cell. n.s., $p > 0.05$; Repeated measures Anova followed by Tukey's multiple comparison test. **(G)** Photomodulation ratios of AMPA-EPSCs from MASp-labeled, control neurons, showing no significant photomodulation. Quantification of the photomodulation ratio was performed similarly to Fig. 3. n.s., $p > 0.05$; one sample Wilcoxon test against the value of 1. **(H-J)** No photodependence of the excitability properties of MASp-labeled control neurons. **(H)** Current-clamp voltage traces of a MASp-labeled, control neuron from a P23 mouse during 500 ms current injection steps under UV (365 nm, violet bar) and green light (525 nm, green bar). UV light was applied 50 ms before and during the current step, while green light was applied 2 s before and during the current step. The amount of current injected is written next to the corresponding voltage trace. **(I,J)** No light-dependence of the membrane potential (I) and firing frequency (J) as a function of injected current after MASp labeling of control neurons from P21-P24 animals. All recordings in brain slices were performed at physiological pH.

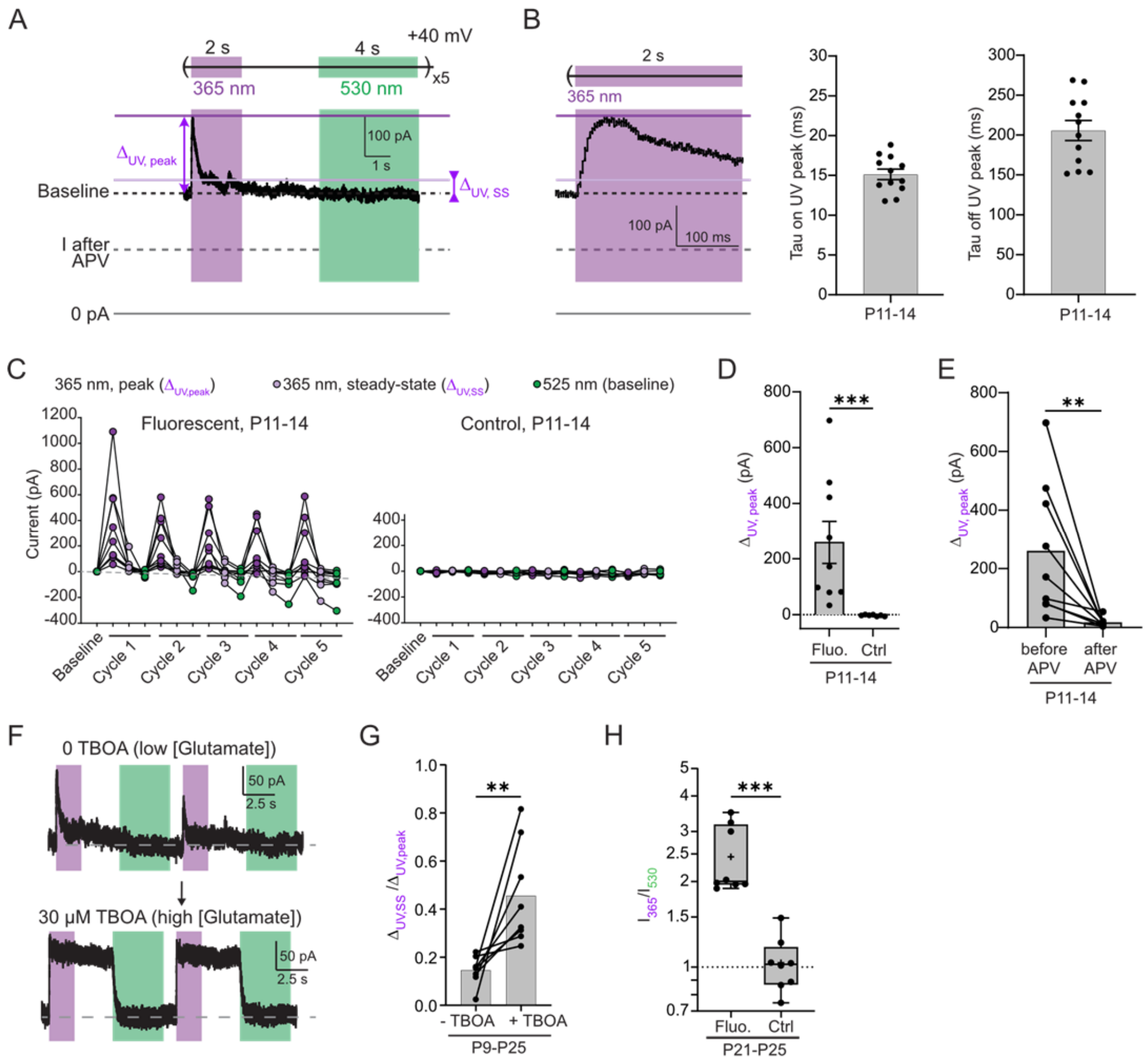


**Fig. S6 (related to Fig. 3):**

Unless otherwise noted, experiments were performed on slices from P11-14 electroporated animals. **(A)** NMDAR tonic current trace of the MASp-labeled, P13 fluorescent neuron shown in Fig. 3D displaying the different values used in the following panels. $\Delta_{UV,\ peak}$ represents the difference between the peak current amplitude at the onset of UV irradiation and the current amplitude under green light (baseline current). $\Delta_{UV,\ SS}$ represents the difference between the steady-state current amplitude at the end of UV irradiation and the current amplitude under green light (baseline current). **(B)** Left, zoom on the UV-induced peak of the NMDA tonic current trace from (A). Middle and right, time constants of UV-induced potentiation (Tau on UV peak, left) and relaxation (Tau off UV peak, right). Tau on UV peak = 15.1 ± 0.7 ms, n = 12 and Tau off UV peak = 206 ± 13 ms, n = 12. **(C)** Tonic current amplitudes of MASp-labeled, fluorescent (left) and control (right) neurons during the 5-cycle illumination protocol. See Panel (A) for annotations. **(D)** UV-induced increase of tonic current amplitude at peak ($\Delta_{UV,\ peak}$) for each MASp-labeled, fluorescent and control (non-fluorescent, denoted ctrl) neuron displayed in Panel C, and averaged over the 5 cycles of illumination. ***, $p < 0.001$, Mann Whitney test. **(E)** APV suppresses UV-induced increase of tonic current at peak of MASp-labeled, fluorescent neurons. **, $p < 0.01$, Mann-Whitney test. **(F)** Photomodulation of the tonic current of a MASp-labeled, P9 fluorescent neuron in basal conditions (0 TBOA, top) and after 10 min treatment of the slice with the glutamate transporter inhibitor TBOA (30 µM TBOA, bottom). As tonic glutamate concentration increases, UV-induced potentiation goes from transient to sustained. **(G)** Relative amplitude of steady-state over peak UV-induced increase in tonic current ($\Delta_{UV, SS}$ / $\Delta_{UV, peak}$) before (-TBOA) and after (+TBOA) TBOA treatment. Animals with ages ranging from P9 to P25 were used for this experiment. **, $p < 0.01$; Wilcoxon matched-pairs signed rank test. **(H)** Photomodulation ratios of NMDA tonic currents of MASp-labeled, fluorescent and control (non-fluorescent) neurons from P21-P25 animals. Photomodulation values and cell numbers are summarized in Table S3. ***, $p < 0.001$, Mann Whitney test. All recordings in brain slices were performed at physiological pH.

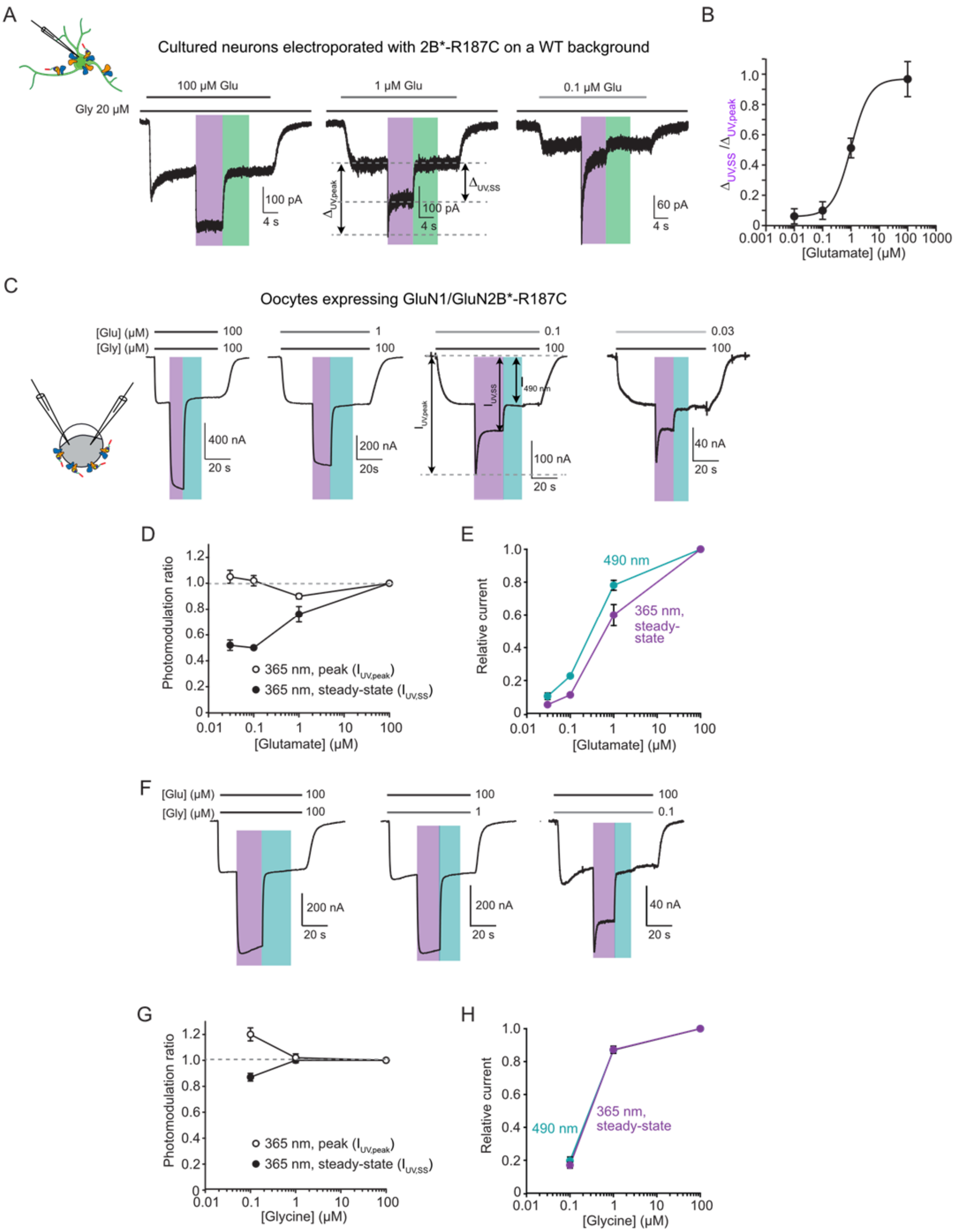


**Fig. S7 (related to Fig. 3): Stability of Opto2B photoenhancement depends on glutamate concentration.**

**(A)** Current traces from a cultured cortical neuron electroporated with the GluN2B*-R187C subunit and labeled with MASp showing transient UV (violet bar)-induced potentiation of NMDAR currents under non-saturating glutamate (Glu) concentrations (0.1 and 1 μM), while UV potentiation was stable at saturating glutamate concentration (100 μM). NMDAR currents were isolated using 10 μM NBQX, 100 μM picrotoxin and 10 μM strychnine, and 20 μM glycine was applied throughout the experiment (same conditions as in *ex vivo* experiments in Fig. 3). The experimental strategy to express the mutated GluN2B subunit on a WT background was the same as in Fig. S4A. **(B)** The ratio of steady-state over peak current ($\Delta_{UV, SS}$ / $\Delta_{UV, peak}$; see Panel (A) for measurement of $\Delta_{UV, SS}$ and $\Delta_{UV, peak}$) increases as glutamate concentration increases. Recordings were performed at pH 7.3, -60 mV. **(C-H)** Experiments were performed in Xenopus oocytes expressing GluN1/GluN2B*-R187C at pH 6.5, -60 mV. **(C-E)** Glutamate dependence of GluN2B diheteromer photomodulation. **(C)** Current traces from an oocyte expressing GluN1/GluN2B*-R187C and labeled with MASp following application of varying concentrations of glutamate together with a saturating concentration (100 μM) of glycine and under 365 (violet bar) or 490 nm (blue-green bar) illumination. **(D)** While the photomodulation ratios at steady-state ($I_{UV, SS}$ / $I_{490\,nm}$, black dots) are highly dependent on glutamate concentration, the photomodulation ratios calculated from the UV peak currents ($I_{UV, peak}$ / $I_{490\,nm}$, white dots) were almost independent of glutamate concentration (they varied by less than 10% across all glutamate concentrations). For each cell, $I_{UV, peak}$ was measured at the time of peak measured for 0.1 μM glutamate, i.e. ~0.9 s after the onset of UV illumination. $I_{UV, SS}$ was measured at the end of the UV illumination step. $I_{490\,nm}$ was measured at the end of the 490 nm illumination step. **(E)** Relative currents at different glutamate concentrations normalized to the current at 100 μM glutamate (saturating concentration). **(F-H)** Glycine dependence of GluN1/GluN2B photomodulation. **(F)** Current traces from an oocyte expressing GluN1/GluN2B*-R187C and labeled with MASp following application of varying concentrations of glycine together with a saturating concentration (100 μM) of glutamate and under 365 (violet bar) or 490 nm (blue-green bar) illumination. **(G)** UV potentiation was stable for 100 and 1 μM glycine but became transient for 0.1 μM glycine. Maximum UV potentiation ($I_{UV, peak}$ / $I_{490\,nm}$, white dots) increased by 20% at low glycine concentrations compared to higher 1 and 100 μM concentrations. **(H)** Relative currents at different glycine concentrations normalized to the current at 100 μM glycine (saturating concentration).

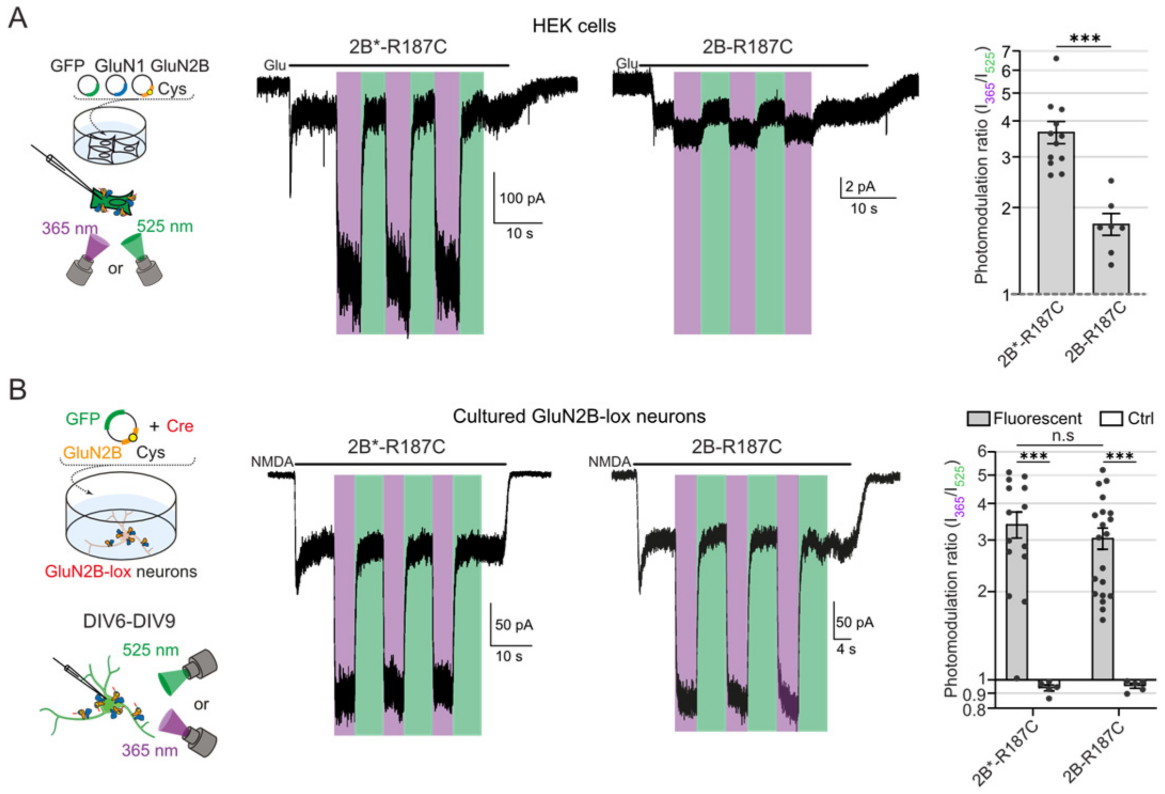


**Fig. S8 (related to Fig. 4): Design strategy of the Opto2B mouse line: neutralizing the endogenous cysteine C395 is not necessary to obtain strong photomodulation in native preparations.**

**(A)** Left, current traces from HEK cells expressing GluN1/GluN2B-R187C-C395S (GluN1/GluN2B*-R187C) and GluN1/GluN2B-R187C receptors (with endogenous cysteine 395 intact), and labeled with MASp, following application of glutamate (100 µM) under 365 nm (violet bars) or 525 nm (green bars) illumination. Cells were constantly perfused with glycine (100 µM). Right, summary of the photomodulation ratios ($I_{365\ nm}$ / $I_{525\ nm}$) for the different GluN2B mutants expressed in HEK cells. Note that the photomodulation ratio is much lower for GluN1/GluN2B-R187C (1.75 ± 0.15, n = 7) compared to GluN1/GluN2B*-R187C (3.66 ± 0.32, n = 12). ***, $p < 0.001$; Mann Whitney test. **(B)** Left, current traces from DIV6-9 cortical neurons expressing GluN1/GluN2B-R187C-C395S (GluN1/GluN2B*-R187C) and GluN1/GluN2B-R187C receptors, and labeled with MASp, following application of NMDA (300 µM) under 365 nm (violet bars) or 525 nm (green bars) illumination. Cells were constantly perfused with D-serine (50 µM). In this particular experiment, we adopted a strategy of molecular replacement of the endogenous, WT GluN2B subunits by the mutated GluN2B subunits using the Cre-lox approach. Dissociated cortical neurons from floxed-Grin2B mice (ref. (1)) were electroporated with pIRES plasmids coding for GFP and a mutated GluN2B subunit (ref. (2)), and plasmids coding for the Cre recombinase (to remove endogenous GluN2B expression) and for the fluorescent marker

TdTomato (see Methods). Right, summary of the photomodulation ratios ($I_{365\ nm}$ / $I_{525\ nm}$) of currents from the different GluN2B mutants expressed in cultured neurons. Note that, contrary to what was observed on HEK cells, the photomodulation ratio is similar between GluN1/GluN2B-R187C (3.04 ± 0.25, n = 20) and GluN1/GluN2B*-R187C (3.25 ± 0.35, n = 15). n.s, $p > 0.05$; ***, $p < 0.001$; Multiple Mann-Whitney tests, p-values were adjusted for multiple comparisons using Bonferroni correction. Only the pre-selected indicated comparisons were performed. All recordings were performed at pH 7.3, -60 mV.

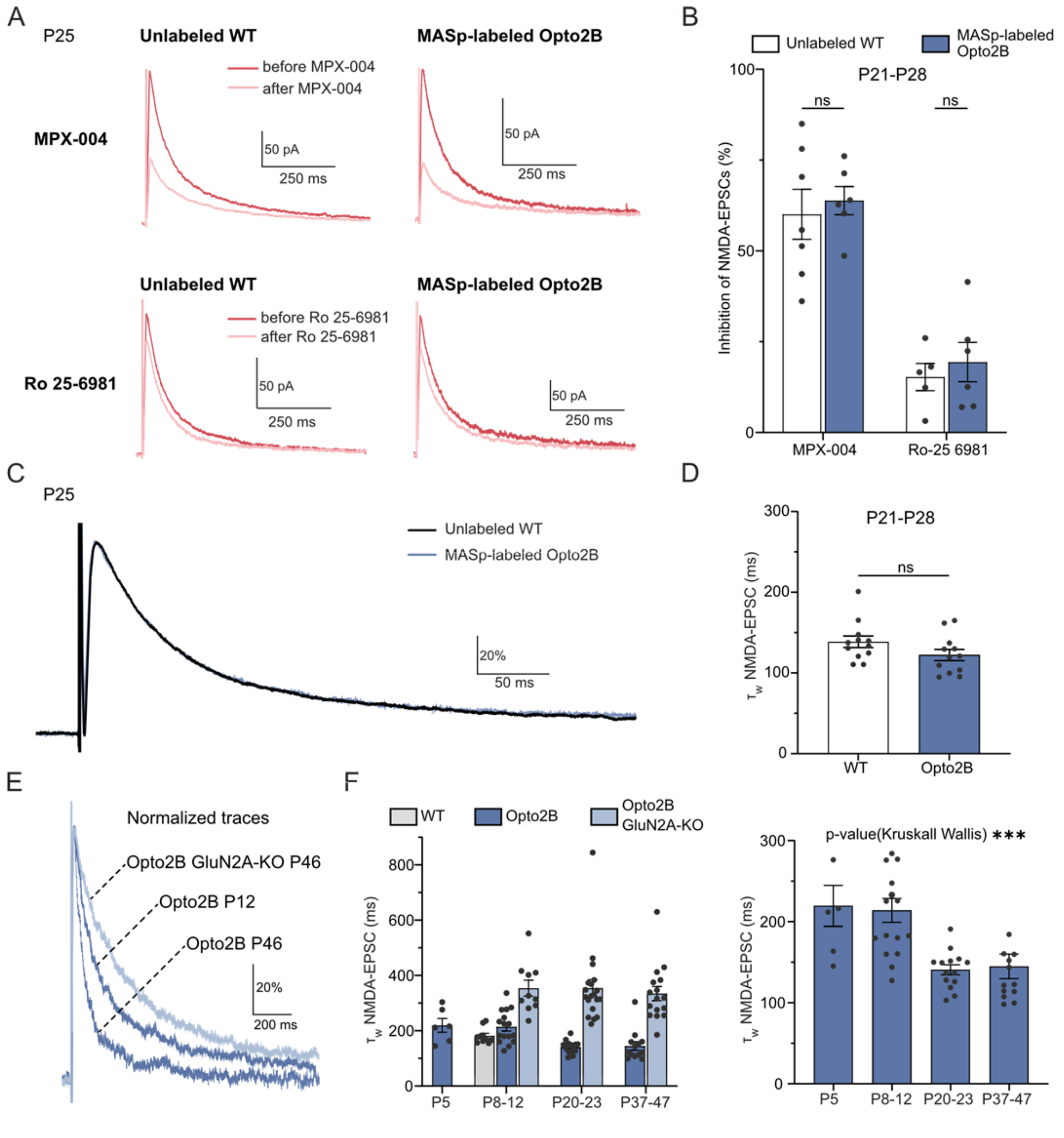


**Fig. S9 (related to Fig. 4): Similar NMDAR subunit profiles between Opto2B and WT mice.** **(A)** Top, superposition of NMDA-EPSCs of CA1 pyramidal neurons before and after application of MPX-004 (top) or Ro 25-6981 (bottom) from unlabeled WT (left) and MASp-labeled Opto2B mice (right) at age P25, recorded in the dark (MASp in its inactive, *trans* configuration). **(B)** Summary of NMDA-EPSC inhibition (%) by MPX-004 or Ro 25-6981 of CA1 pyramidal neurons from unlabeled WT (white) and MASp-labeled Opto2B mice (blue) at P21-P28. n.s., $p > 0.05$; multiple Mann-Whitney tests, p-values were adjusted using Bonferroni correction. Only the pre-selected indicated comparisons were performed. **(C)** Superposition of normalized NMDA-EPSCs of CA1 pyramidal neurons from unlabeled WT (black) and MASp-labeled Opto2B mice (blue) at age P25. **(D)** Summary of NMDA-EPSC decay kinetics from unlabeled WT (white) and MASp-labeled

Opto2B mice (blue) at P21-P28. n.s., $p > 0.05$; Mann Whitney test. Note the similar NMDA-EPSC decay time between WT and Opto2B animals, suggesting that the R187C mutation does not perturb developmental maturation of NMDA subtypes. **(E)** Superposition of normalized NMDA EPSCs of MASp-labeled CA1 pyramidal neurons from Opto2B mice at age P12 and P46, as well as from P46 Opto2B/GluN2A KO animals, recorded under green light (MASp in its inactive configuration). Each trace is the average of 12 individual EPSCs. **(F)** Left, decay time constants ($\tau_w$ NMDA-EPSC) of NMDA-EPSCs of MASp-labeled CA1 pyramidal neurons from WT (grey), Opto2B (dark blue) and Opto2B GluN2A-KO (light blue) mice at different age ranges. Note the absence of significant difference between the NMDA-EPSC decay kinetics of from WT and Opto2B animals at P8-P12, suggesting similar NMDAR subunit populations. NMDA-EPSC decay kinetics from Opto2B/GluN2A KO mice remained slow at older ages, consistently with what was previously shown for animals lacking GluN2A expression (ref. (3)). Right, zoom on the Opto2B mouse condition for better visualization. ***, $p < 0.001$, Kruskall-Wallis test on Opto2B condition. Note the acceleration of NMDA-EPSC decay time with age on Opto2B mice, which is consistent with what was observed in the literature on WT animals (ref (4)). All the recordings on brain slices were performed at physiological pH.

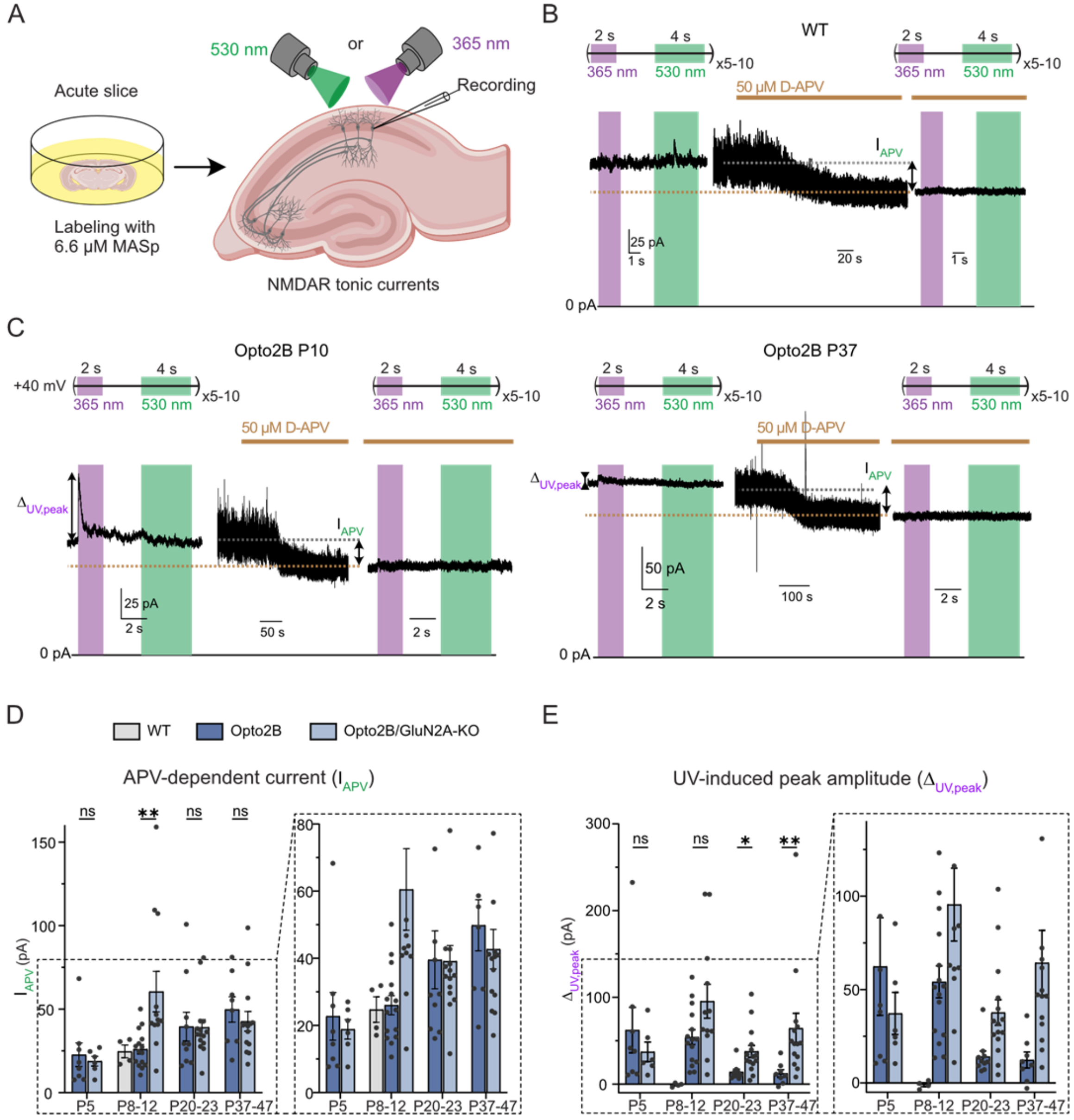


**Fig. S10 (related to Fig. 5):**

**(A)** Protocol of tonic current photomodulation. Note the absence of electrical stimulation: recorded tonic currents are mediated by low levels of tonic glutamate. See Methods and Main Text for more details. **(B)** Tonic current trace of a MASp-labeled, CA1 pyramidal neuron from a P10 WT mouse before (left) and during (middle and right) application of 50 μM D-APV. Left and right traces are the average of 5 to 10 traces. Note the absence of photomodulation. **(C)** Tonic current traces of MASp-labeled, CA1 pyramidal neurons from a P10 (left) or a P37 (right) Opto2B mouse before and during application of 50 μM D-APV. Left and right traces are the average of 5 to 10 traces. Displayed arrows indicate the currents measured in the following panels and used to calculate the photomodulation ratio (see also Fig. 3). **(D)** Amplitude of APV-sensitive currents (as shown in a) of MASp-labeled, CA1 pyramidal neurons from WT (grey), Opto2B (dark blue) and Opto2B GluN2A-KO (light blue) mice according to age ranges. Right, zoom on data for better visibility of bar graphs. **(E)** Amplitude of the UV-induced peak (as shown in a) of MASp-labeled, CA1 pyramidal neurons from WT (grey), Opto2B (dark blue) and Opto2B GluN2A-KO (light blue) mice according to age ranges. Right, zoom on data for better visibility of bar graphs. All recordings in brain slices were performed at physiological pH.

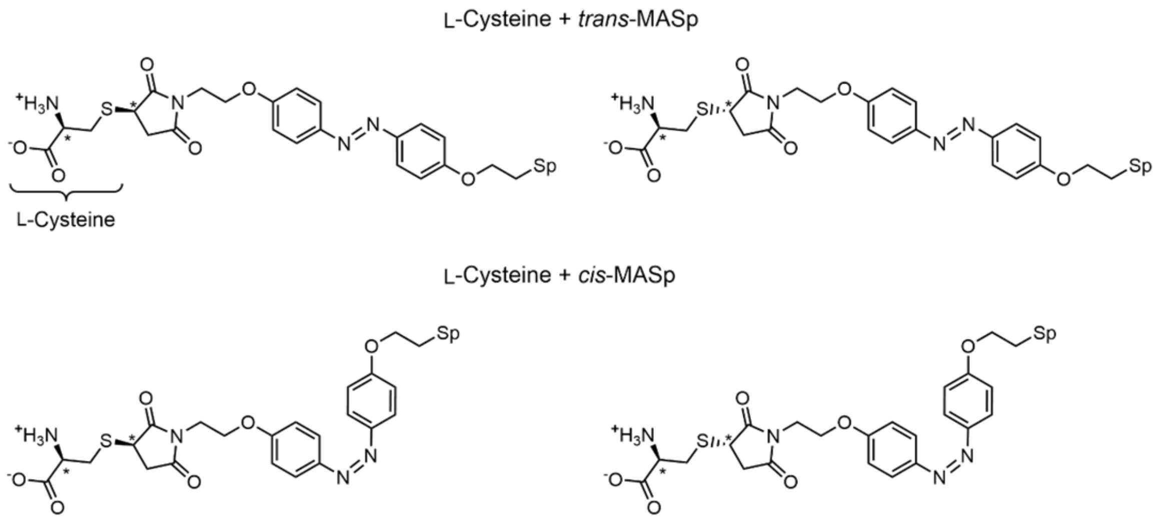


**Fig. S11: Chemical structures of the four diastereoisomeric products of the reaction between MASp and L-Cysteine.** Stars indicate asymmetric carbons.

#### Supplementary Tables

**Table S1: Summary of *in vitro* photomodulation data.** Photomodulation steady-state values of different NMDAR constructs labeled with MASp in Xenopus oocytes, HEK cells and cultured cortical neurons.

| Construct | $I_{UV} / I_{green}$ [a] | | | |
|---|---|---|---|---|
| | **Oocytes (pH 6.5)** | **Oocytes (pH 7.3)** | **HEK cells (pH 7.3)** | **Neurons (pH 7.3)** |
| **2B WT** | 1.00 ± 0.02 (n = 20) | 0.99 ± 0.01 (n = 4) | 1.048 ± 0.005 (n = 5) | - 0.968 ± 0.009 (n = 9) (non-fluorescent neurons on WT background; Fig. S4)<br><br>- 0.95 ± 0.02 (n = 5) (2B*-R187C condition) 0.94 ± 0.02 (n = 5) (2B-R187C condition) (non-fluorescent neurons ; molecular replacement strategy ; Fig. S8B) |
| **2B*-Q180C** | 0.39 ± 0.02 (n = 27) | 0.84 ± 0.08 (n = 13) | 0.77 ± 0.07 (n = 4) | |
| **2B*-R187C** | 3.53 ± 0.21 (n = 32) | 1.97 ± 0.17 (n = 21) | 2.9 ± 0.7 (n = 8) (Fig. S3)<br><br>3.7 ± 0.3 (n = 12) (Fig. S8a) | - 1.81 ± 0.09 (n = 19) (fluorescent neurons on WT background; Fig. S4)<br><br>- 3.40 ± 0.35 (n = 14) (fluorescent neurons ; molecular replacement strategy ; Fig. S8B) |
| **2B-R187C** | | | 1.75 ± 0.40 (n = 7) | - 3.04 ± 0.25 (n = 20) (fluorescent neurons ; Cre-lox strategy) |
| **2A WT** | 0.92 ± 0.02 (n = 22) | | 0.88 ± 0.02 (n = 4) | |
| **2C WT** | 1.02 ± 0.01 (n = 5) | | | |
| **2D WT** | 1.00 ± 0.01 (n = 5) | | | |
| **2B*-R187C-r1 / 2B*-R187C-r2** | 3.07 ± 0.22 (n = 9) | | | |
| **2B-r1 / 2B*-R187C-r2** | 1.21 ± 0.02 (n = 14) | | | |
| **2A-r1 / 2B*-R187C-r2** | 1.01 ± 0.01 (n = 17) | | | |

[a] Green light is 490 nm for oocytes and 525 nm for HEK cells and neurons.

**Table S2**: **Summary of several pharmacological parameters of 2B*-Q180C and 2B*-R187C mutants unlabeled (- MASp) and labeled (+ MASp) with MASp under different illumination conditions.**

| | | Glu $EC_{50}$ | Gly $EC_{50}$ | Relative Po[a] | $I_{pH\ 7.3}$ / $I_{pH\ 6.5}$ | $I_{spermine}$ / $I_0$[b] |
|---|---|---|---|---|---|---|
| **2B wt** | **- MASp** | 1.44 ± 0.15 (n = 9) | 0.48 ± 0.09 (n = 4) | 1.01 ± 0.04 (n = 15) | 10.4 ± 0.3 (n = 20) | 8.0 ± 0.4 (n = 21) |
| **2B*-Q180C** | **- MASp** | 1.43 ± 0.10 (n = 3) | 0.29 ± 0.03 (n = 5) | 0.96 ± 0.1 (n = 8) | 9.5 ± 0.6 (n = 11) | 7.5 ± 0.5 (n = 11) |
| | **+ MASp, 365 nm** | 1.83 ± 0.12 (n = 7) | 0.29 ± 0.06 (n = 3) | 0.76 ± 0.14 (n = 7) | 9.5 ± 0.8 (n = 6) | 4.3 ± 0.4 (n = 9) |
| | **+ MASp, 490 nm** | 2.49 ± 0.11 (n = 6) | 0.57 ± 0.08 (n = 6) | 2.15 ± 0.04 (n = 9) | 5.2 ± 0.3 (n = 5) | 2.7 ± 0.2 (n = 8) |
| **2B*-R187C** | **- MASp** | 0.94 ± 0.04 (n = 11) | 0.49 ± 0.05 (n = 8) | 0.64 ± 0.04 (n = 6) | 13.4 ± 0.6 (n = 9) | 11.5 ± 0.8 (n = 9) |
| | **+ MASp, 365 nm** | 1.83 ± 0.08 (n = 7) | 0.30 ± 0.03 (n = 4) | 2.22 ± 0.19 (n = 5) | 6.9 ± 0.5 (n = 7) | 4.3 ± 0.4 (n = 7) |
| | **+ MASp, 490 nm** | 0.75 ± 0.07 (n = 8) | 0.43 ± 0.05 (n = 4) | 0.56 ± 0.04 (n = 5) | 9.5 ± 0.5 (n = 7) | 7.5 ± 0.7 (n = 7) |

[a]Relative Po is measured as the rate of MK-801 inhibition normalized to the average rate of inhibition of wt GluN2B-NMDARs measured the same experimental day.
[b]Spermine was applied at a concentration of 200 µM (around its $EC_{50}$, see ref. (5)).
Cell numbers are indicated in parenthesis. Glu, glutamate; Gly, glycine.

**Supplementary Table 3: Summary of *ex vivo* photomodulation data on cortical neurons**

| Photomodulation ($I_{365\ nm}$ / $I_{530\ nm}$) | | Opto2B neurons | Control neurons |
|---|---|---|---|
| **P11-14** | **NMDA EPSCs** | 1.45 ± 0.04 (n = 9) | 0.99 ± 0.02 (n = 7) |
| | **NMDA tonic current** | 3.83 ± 0.45 (n = 9) | 0.95 ± 0.03 (n = 7) |
| **P21-25** | **NMDA EPSCs** | 1.27 ± 0.03 (n = 9) | 1.01 ± 0.02 (n = 7) |
| | **NMDA tonic current** | 2.44 ± 0.24 (n = 8) | 1.04 ± 0.08 (n = 8) |

**Supplementary Table 4: Summary of *ex vivo* photomodulation data in hippocampal CA1 pyramidal cells**

| Photomodulation ($I_{365\ nm}$ / $I_{530\ nm}$) | | WT | Opto2B | Opto2B / GluN2A KO |
|---|---|---|---|---|
| **P5** | **NMDA EPSCs** | | 1.56 ± 0.17 (n = 6) | |
| | **NMDA tonic current** | | 3.53 ± 0.55 (n = 8) | 3.05 ± 0.47 (n = 6) |
| **P8-12** | **NMDA EPSCs** | 1.02 ± 0.04 (n = 10) | 1.41 ± 0.06 (n = 20) | 2.07 ± 0.20 (n = 10) |
| | **NMDA tonic current** | 0.98 ± 0.04 (n = 4) | 3.30 ± 0.39 (n = 15) | 2.53 ± 0.13 (n = 12) |
| **P20-23** | **NMDA EPSCs** | | 1.28 ± 0.04 (n = 14) | 1.63 ± 0.06 (n = 22) |
| | **NMDA tonic current** | | 1.46 ± 0.08 (n = 10) | 2.12 ± 0.20 (n = 15) |
| **P37-47** | **NMDA EPSCs** | | 1.14 ± 0.03 (n = 14) | 1.51 ± 0.05 (n = 17) |
| | **NMDA tonic current** | | 1.32 ± 0.12 (n = 8) | 2.67 ± 0.42 (n = 14) |

**Supplementary Table 5 (separate file): Summary of all statistical tests**
